# Functional connectivity is linked to symbolic BOLD patterns: replication, extension, and clinical application of the human ‘complexome’

**DOI:** 10.1101/2025.09.05.674447

**Authors:** Amy Romanello, Nina von Schwanenflug, Friedemann Paul, Harald Prüss, Stephan Krohn, Carsten Finke

**Affiliations:** Department of Neurology and Experimental Neurology, Charité-Universitätsmedizin Berlin, corporate member of Freie Universität Berlin, Humboldt-Universität Berlin, Berlin, Germany; Berlin School of Mind and Brain, Humboldt-Universität zu Berlin, Berlin, Germany; Max Delbrück Center for Molecular Medicine in the Helmholtz Association, Berlin, Germany; Experimental and Clinical Research Center, a cooperation between the Max Delbrück Center for Molecular Medicine in the Helmholtz Association and Charité—Universitätsmedizin Berlin, Berlin, Germany; NeuroCure Clinical Research Center, Charité—Universitätsmedizin Berlin, corporate member of Freie Universität Berlin, Humboldt-Universität zu Berlin, and Berlin Institute of Health, Berlin, Germany; German Center for Neurodegenerative Diseases (DZNE) Berlin, Berlin, Germany

## Abstract

Functional connectivity (FC) quantifies the temporal coherence of blood-oxygen-level-dependent (BOLD) signals across brain regions. Recently, the information-theoretic “complexome” framework has linked FC to coinciding ‘complexity drops’: transient moments in which regional BOLD signals simultaneously become regular. Here, we replicate this relationship in an independent dataset and extend the framework by (i) integrating it with signal cofluctuation analysis through edge-timeseries, (ii) extending the previous binary concept of simultaneous complexity drops to a continuous, threshold-free calculation, (iii) providing evidence of clinical relevance in the model disease of anti-N-methyl-D-aspartate-receptor encephalitis, and (iv) deriving a novel measure of pairwise dissimilarity in local BOLD patterns. This ‘index of pattern incongruency’ (IPI) explains clinically relevant FC reductions and maps onto novel associations with cognition. These findings show that global FC is closely related to local patterns within underlying BOLD signals, strengthening the link between complexity dynamics and the brain’s functional organization as a large-scale network.

## Introduction

Neural variability is a fundamental feature of human brain function (*1*, *2*). Functional magnetic resonance imaging (fMRI) has been instrumental in advancing our understanding of these brain dynamics, both in health and across various clinical conditions. A central approach in human neuroimaging is the estimation of functional connectivity (FC) from blood-oxygen-level-dependent (BOLD) signals in resting-state fMRI (*3*). FC refers to the covariance structure of BOLD signals across the brain, where two brain regions are considered functionally connected if their signals show a high degree of temporal coherence – commonly estimated as the product-moment correlation of the two signal vectors. Within this framework, the brain is viewed as a large-scale functional network, in which gray matter regions represent the *nodes* of the network, and the FC estimates represent the *edges* between these nodes.

While this approach has been widely adopted in neuroimaging research, efforts to establish a clear mechanistic link between the brain’s global network architecture and the local neural dynamics from which it emerges are comparatively recent. Elucidating this relationship is crucial, since the estimation of FC as temporal correlation represents a non-injective function. That is, although a pair of BOLD signals uniquely define a specific FC value, many other signals can produce the same value, even if their underlying signal properties fundamentally differ. In consequence, one can reliably infer FC from neural dynamics, but not vice versa: the neural dynamics of two individual regions cannot be uniquely reconstructed from their FC. Therefore, a complete understanding of the brain’s functional architecture requires a link between node-level activity and edge-level connectivity of the network.

Recently, progress in this regard has been made by two complementary lines of research, which can be subsumed as an *edge-centric* and a *node-centric* account of brain activity, respectively.

On the one hand, Esfahlani et al. (2020) introduced the so-called ‘edge-timeseries’ framework. This approach rests on a continuous measure of BOLD signal cofluctuation that captures bidirectional contributions to the FC of any given set of regions at the single time-point resolution. Empirical findings suggest that brief moments with highest amplitude of BOLD cofluctuation drive FC strength and relate to the canonical network architecture observed in resting-state brain activity (*4*).

On the other hand, we recently presented the ‘complexome’ framework – an information-theoretic account of nodal activity, which is rooted in the encoding of symbolic patterns within local BOLD signals (*1*). Specifically, we used a time-resolved computation of weighted permutation entropy (WPE) (*5*) to estimate the complexity of regional BOLD signals from the distribution of these patterns over a given moment in time. In this framework, a low-complexity BOLD signal is one where a dominant pattern explains most of the amplitude variance (i.e., the signal is more regular), whereas a high-complexity signal features various symbolic patterns that split the amplitude information between them (i.e., the signal is more irregular). With this approach, we found that the brain operates in a default state of high complexity over most of the resting-state recording. However, this default activity is repeatedly interrupted by transient moments of increased signal regularity, which become visible as spontaneous ‘complexity drops’. Importantly, we previously showed that these complexity dynamics cannot be explained by signal covariance alone and are distinct from BOLD cofluctuations quantified with the original edge-timeseries approach (*1, see fig. S6*). Furthermore, we observed that the FC strength between any two brain regions is closely related to how often they engage in these complexity drops simultaneously (i.e., their ‘drop coincidence’). This finding establishes a systematic link between local complexity dynamics (node level) and interregional FC (edge level).

Given their unique yet complementary contributions, integrating these frameworks is an important next step in understanding the link between local activity and global connectivity of the brain. By combining both approaches, we extend the ‘complexome’ framework to *complexity cofluctuations*, creating a unified model that integrates the global properties of FC, the complexity dynamics of regional BOLD signals, and the abstract patterns within those signals. This integration yields several methodological extensions, enhancing its applicability to an even wider range of research questions. Specifically, the original definition of drop coincidence rested on a binary drop threshold—the first percentile of empirically observed WPE values. However, this definition excludes moments in which two regions exhibit simultaneous decreases in complexity, but only one meets the drop criterion. Borrowing from the mathematical formulation of edge-timeseries, the concept of drop coincidence can be extended to a continuous, threshold-free calculation that also captures more subtle occasions of simultaneous complexity decrease. Moreover, this combined approach offers a more granular account of BOLD signal dynamics by considering the specific pattern distributions that underlie both FC and WPE. As such, it provides a powerful tool for studying hierarchical brain dynamics and for investigating their relevance for cognition and clinical symptoms. This is particularly important in neurological populations where FC analyses have been crucial in understanding pathological brain changes, e.g., associations with structural atrophy and network disruptions in conditions such as dementia (*6*), multiple sclerosis (*7*), and autoimmune encephalitis (*8*, *9*). Yet, these FC alterations may result from many different signal fluctuations, making the neurobiological interpretation of such findings a challenge. Clarifying how local signal patterns relate to global connectivity is therefore a crucial step towards clinical translation and the generation of more mechanistic insights into neurological disease.

Against this background, the aims of our study were to (i) leverage the recent development of edge-timeseries and transfer it to complexity dynamics (‘complexity cofluctuations’), (ii) extend the previous binary concept of drop coincidence to a continuous, threshold-free calculation, (iii) derive a new measure of pairwise dissimilarity in local BOLD signal patterns, (iv) apply the complexome framework to an independent dataset and scanning protocol, and (v) test the sensitivity of the framework to clinically relevant FC changes. To this end, we study both healthy control participants (HC) and a large cohort of patients with anti-N-methyl-D-aspartate receptor (NMDAR) encephalitis, the most common form of autoimmune encephalitis (*10*). Patients with NMDAR encephalitis exhibit a severe neuropsychiatric syndrome associated with altered brain dynamics in fMRI (*11–13*), specifically reduced FC in hippocampal connections with the default mode network (DMN) (*9*, *12*, *14*, *15*). Given the robustness and replicability of this finding, NMDAR encephalitis here serves as a ‘model disease’ to explore if complexity dynamics can explain well-established and clinically relevant changes in FC.

## Results

### Extending the ‘complexome’: complexity cofluctuations and pattern incongruency

Time-resolved BOLD signal complexity was computed using a sliding window approach, as previously described in (*1*) and detailed in the Materials and Methods. This yielded regional complexity timeseries at the resolution of temporal windows (Fig. 1a). Subsequently, we adapted the recent edge-timeseries framework of Esfahlani et al. (2020) to compute the pairwise cofluctuation between these complexity timeseries (Fig. 1b, left). This approach allowed us to extend the binary concept of drop coincidence to a continuous, threshold-free metric: specifically, we take the root-sum-square of the element-wise product between complexity timeseries at instances of simultaneous complexity decreases (i.e., downward cofluctuations) and summarize them as the area-under-the-curve (AUC) over time. A decrease in complexity was defined relative to a signal’s complexity value during the previous window. Thus, this measure captures both the magnitude (amplitude of cofluctuation) and duration (number of windows) of simultaneous complexity decrease (SCD) between a given pair of brain regions (Fig. 1b, right). Furthermore, we examined the individual BOLD signal patterns during these windows of SCD. Using the pattern frequency distributions produced during the calculation of WPE, we computed the Euclidean distance between a given pair of distributions in every window that showed a SCD. Then, we defined an index of pattern incongruency (IPI) as the mean distance over all these windows (Fig. 1c). IPI thus serves as a summary measure of pairwise BOLD signal complexity dynamics that relates underlying nodal activity patterns to the strength of edgewise functional coupling.

**Figure 1.**
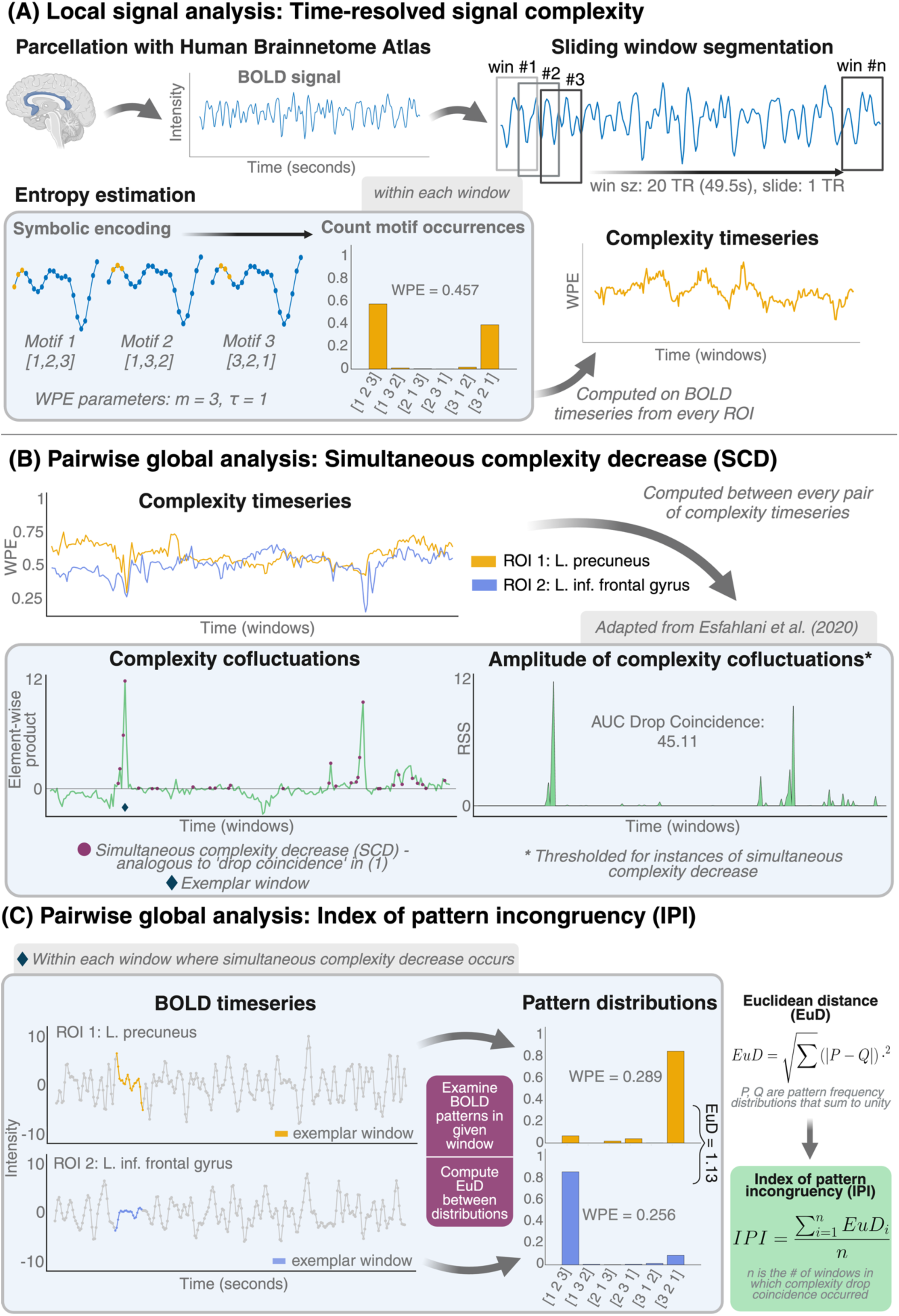
Computing time-resolved signal complexity and BOLD pattern incongruency. **(A)** The step-by-step process for computing weighted permutation entropy (WPE). Preprocessed fMRI data was parcellated into 244 regions of interest using the Human Brainnetome Atlas (*16*). The sliding window technique was used to segment regional timeseries into consecutive windows of 20 TRs (49.5s), with an overlap of 95%. Within each window, WPE was calculated using a rank-based symbolic encoding framework depicted in the blue inset panel (see Materials and Methods). This resulted in a set of 244 complexity timeseries per participant at the resolution of windows. **(B)** Transferring edge-timeseries to complexity dynamics. A pair of exemplar complexity timeseries is shown on the top-left. The element-wise product between the two z-scored complexity timeseries results in a third timeseries that captures the magnitude and direction of complexity cofluctuations. Positive values here indicate windows in which the complexity timeseries fluctuate in the same direction, while negative values indicate windows with fluctuations in opposing directions. Purple circles indicate windows in which both complexity timeseries showed downward fluctuations, i.e., both underlying BOLD timeseries showed increased regularity compared to the previous window. The complexity cofluctuation timeseries was indexed to retain amplitudes corresponding to simultaneous downward cofluctuation. The magnitude of these downward cofluctuations was calculated as the root-sum-square (right) and summarized as the area-under-the-curve (AUC) over time, yielding a continuous, threshold-free measure of simultaneous complexity decrease (SCD). **(C)** The step-by-step process for computing the novel index of pattern incongruency (IPI). The plots show the two BOLD timeseries underlying the exemplar complexity timeseries from panel (B) as well as the corresponding amplitude-weighted pattern frequency distributions. The Euclidean distance is taken between this pair of frequency distributions to compute the pattern dissimilarity over this temporal window. This process is repeated for every window in which simultaneous downward cofluctuations in complexity occurred. The IPI is then defined as the mean over all window-wise Euclidean distances between a pair of BOLD signals (right). Given the mathematical formulation of Euclidean distance (panel C), IPI can take on values between 0 and ✓2. An IPI of exactly 0 indicates that a pair of BOLD signals have identical pattern frequency distributions. Consequently, an IPI of exactly ✓2 indicates that a pair of BOLD signals have maximally distinct pattern frequency distributions.

### Functional connectivity is linked to complexity dynamics and BOLD signal patterns

FC was predominantly positive across the brain in our normative sample of healthy controls (HC). The strongest FC was observed as intra-regional clusters, e.g., with the occipital cortex, pre-central gyrus, and thalamus. Weak anti-correlations were observed mainly between subcortical regions (thalamus, basal ganglia) and primary networks (visual, somatomotor), as well as with task-positive networks (dorsal and ventral attention, frontoparietal) (Fig. 2a, left).

**Figure 2.**
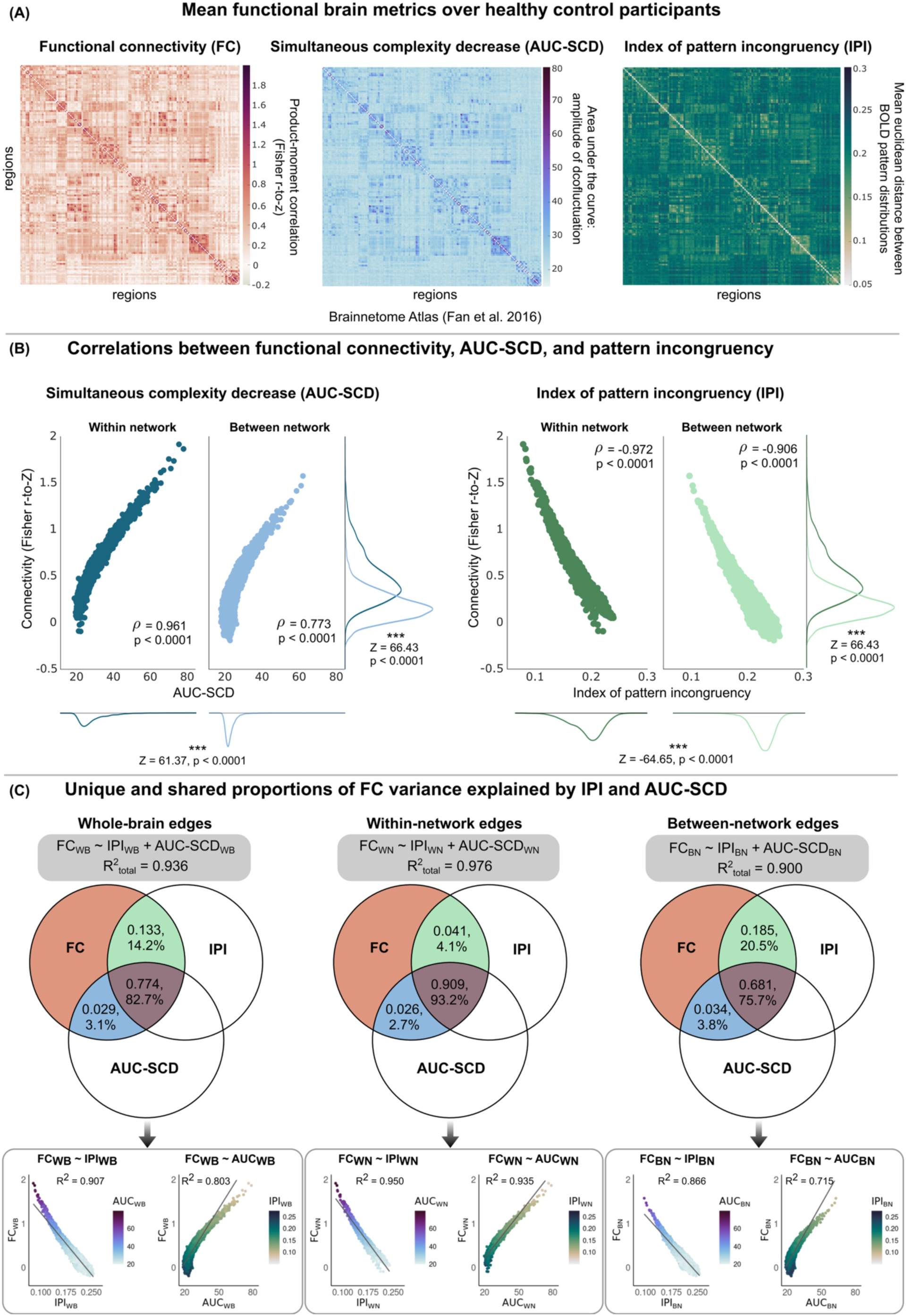
Associations between functional connectivity, AUC-SCD, and BOLD pattern incongruency in healthy participants. **(A)** (Left) Fisher R-to-Z transformed product-moment correlation coefficient. (Middle) Simultaneous complexity decrease (SCD) (Area-under-the-curve, AUC). (Right) Index of pattern incongruency (IPI): Euclidean distance between pairwise BOLD pattern distributions. **(B)** (Left) FC and AUC-SCD show strong positive associations, especially along within-network connections. (Right) FC and IPI show strong inverse associations, especially along within-network connections. Test statistics were calculated using the Wilcoxon rank-sum test. Correlations were computed using the non-parametric Spearman’s rho test. **(C)** Variance in FC across the whole brain as well as within-network and between-network edges is explained by IPI and AUC-SCD. Formulas and the overall coefficient of determination (R^2^) for each multiple regression model are shown for each analysis in the grey boxes. Venn diagrams show the proportion of explained variance in FC that is shared between IPI and AUC-SCD (center, brown), unique to IPI (top, green), and unique to AUC-SCD (left, blue). Unique and shared proportions of variance were calculated using commonality analysis to calculate commonality coefficients according to (*19*). The bottom scatter plots show the individual simple linear regression (SLR) models that were also used in commonality analyses. Color maps of the scatter plots correspond to the second independent variable not included in the individual SLR models. *WB: whole-brain; WN: within-network; BN: between-network*.

FC was strongly linked to AUC-SCD across the whole brain (ρ = 0.82, p < 0.0001), closely replicating findings from our original work (*1*). Similarly, we found that the spatial distribution of signal complexity across the brain was very similar between our in-house HC participants and those from the Human Connectome Project (HCP) Young Adults dataset analyzed in our previous work (r = 0.87, p < 0.0001; Fig. S1), despite differences in data acquisition protocols (*17*). Moreover, SCD were observed across all brain regions, such that every pair of brain regions had a non-zero AUC in every scan. Occipital and frontal regions, as well as the cingulate gyri, showed the greatest AUC-SCD (Fig. 2a, middle). Even with this extended concept of complexity drop coincidence, we found a significant convergence between the spatial distribution of mean AUC-SCD in our in-house HC participants and the binary complexity drop coincidence in the HCP dataset (r = 0.65, p < 0.0001).

Given that we closely replicated the strong link between FC and complexity dynamics in an independent dataset, we next hypothesized that FC should also be related to the BOLD pattern distributions that underlie the WPE calculation. With our IPI measure, we aimed to establish a link between the FC strength of a pair of brain regions and the frequency of particular patterns in the BOLD signals underlying that covariance. Indeed, we observed a strong anti-correlation between FC and IPI across the whole brain (ρ = −0.92, p ∼ 0) – that is, two regions show high temporal covariance (i.e., high FC) if their signals contain very similar patterns (i.e., low IPI). The spatial distribution of IPI across the brain followed an inverse relationship with that of AUC-SCD (ρ = −0.76, p < 0.0001), showing the highest dissimilarities between subcortical regions (namely, the thalamus and basal ganglia) and regions of the somatomotor network as well as the orbital gyri, inferior frontal, and insular regions (Fig 2a, right).

Furthermore, we examined the relationship between FC, AUC-SCD, and IPI over edges *within* and *across* canonical resting-state networks (RSNs). Regional signals were categorized into RSNs according to (*18*) and included the default mode network (DMN), dorsal (DAN) and ventral attention networks (VAN), the frontoparietal network (FPN), limbic network (LIM), somatomotor network (SMN), and the visual network (VIS). Additionally, regions of the left and right hippocampus, amygdala, thalamus, and basal ganglia were included. Within-network connections refer to those edges between regions classified into the same network or subcortical region, while across-network connections refer to edges between two distinct networks or subcortical regions. As expected from the original complexome study (*1*), both FC and AUC-SCD were significantly higher in edges within networks compared to those between networks (FC: Z = 66.43, p < 0.0001; AUC-SCD: Z = 61.37, p < 0.0001). On the contrary, IPI was significantly higher in *between*-network connections (Z = −64.65, p < 0.0001) – that is, neural activity in regions belonging to different functional networks more often show *incongruent* BOLD patterns (Fig. 2b). The relationship between connectivity and AUC-SCD was consistently positive while the relationship between connectivity and IPI was inverse (Fig. 2b). Notably, while both measures were related to within-network FC to a similar degree (Δ|ρ| = 0.01), pattern incongruency was more strongly related to FC for between-network connections (Δ|ρ| = 0.13).

### IPI and AUC-SCD capture unique information about FC

Using multiple regression and commonality analyses (*19*), we observed that the variance in FC strength across the brain is largely explained by IPI and AUC-SCD (Fig. 2c). Since IPI and AUC-SCD themselves are related, we found that the majority of variance in FC is explained by the proportion of variance shared by both variables. However, we also found that both IPI and AUC-SCD exhibit unique contributions to the explanation of FC, evidenced by the commonality coefficients displayed within the Venn diagrams (Fig. 2c). Interestingly, in the case of between-network connections, IPI alone contributes over 20% of the variance explained in FC, highlighting that this novel metric captures unique information about functional dynamics in parallel to AUC-SCD.

In sum, the link between FC and complexity dynamics closely replicates in an independent dataset with a different scanning protocol: regions with high FC more often engage in simultaneous decreases in complexity (AUC-SCD), and this AUC-SCD strongly distinguishes connections within the same functional network from those across different functional networks. Moreover, extending the framework to account for pattern congruency yielded significant additional information: regions show high FC if the underlying BOLD signals contain similar symbolic patterns (i.e. low IPI). This approach tracked global FC even more strongly, which was mostly driven by connections between canonical resting-state networks. Given these strong links in the healthy brain, we next asked if this relationship would also hold in patients with well-established FC alterations.

### Clinical application

Overall, patients with NMDAR encephalitis showed very similar associations between FC, AUC-SCD and IPI (Fig. 3): brain regions with high FC tend to show complexity decreases at the same time. Furthermore, in these moments of SCD (i.e., both BOLD signals are more regular), the signal patterns in both regions are similar to one another (i.e., low IPI). Furthermore, IPI and AUC-SCD captured both shared and unique proportions of variance in FC. Patients also exhibited stronger associations between brain measures in within-network edges compared to those between networks.

**Figure 3.**
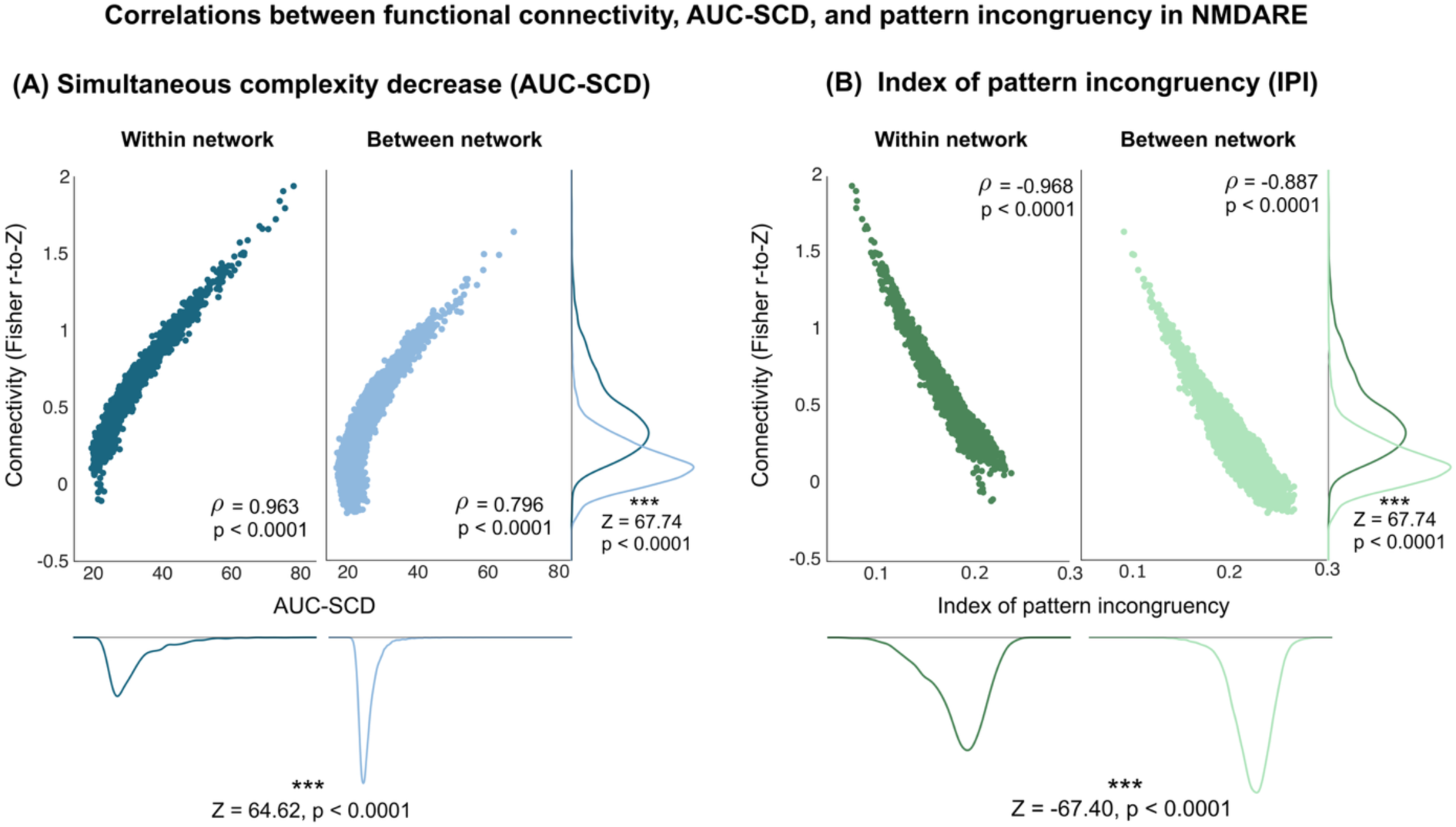
Associations between functional connectivity, AUC-SCD, and BOLD pattern incongruency in patients with NMDAR encephalitis. Patients exhibit very similar relationships across global brain metrics compared to healthy participants. (A) FC and AUC-SCD show strong positive associations, especially along within-network connections. (B) FC and IPI show strong inverse associations, especially along within-network connections. Test statistics were calculated using the Wilcoxon rank-sum Z-test. Correlations were computed using the non-parametric Spearman’s rho test.

These findings demonstrate a consistent relationship between FC, AUC-SCD, and BOLD IPI –both in normative data and in a clinical sample. Therefore, we next studied if disease-related *changes* in FC could be equally well explained by complexity dynamics.

### Hippocampal connectivity alterations in patients with NMDAR encephalitis

Previous studies have consistently observed FC reductions of the hippocampus in patients with NMDAR encephalitis (*9*, *11*, *14*). Therefore, we aimed to (i) replicate these hippocampal FC changes and (ii) investigate to which extent these changes could be explained by the pattern incongruency of the underlying BOLD signals.

Hippocampal functional connectivity analysis confirmed reduced connectivity in patients with NMDAR encephalitis, specifically between bilateral hippocampi and several regions of the default mode network (DMN), including the medial prefrontal cortex (mPFC), orbital gyri, and anterior and posterior cingulate gyri (Fig. 4a, left). Patients also exhibited reduced FC between the bilateral hippocampi and the thalamus (Table S1). There were no hippocampal connections in which patients showed increased FC compared to HC.

**Figure 4.**
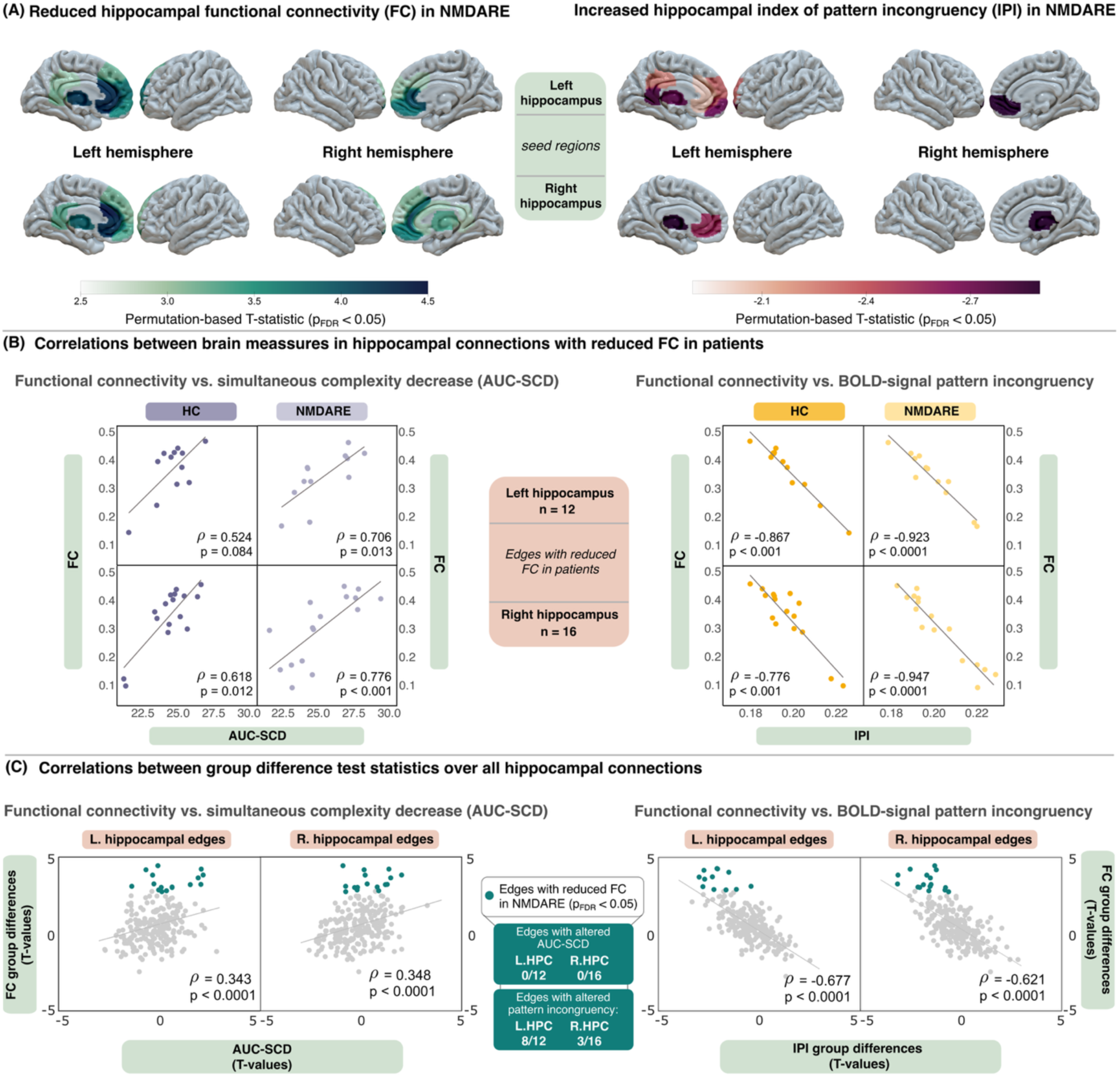
Hippocampal FC reductions in NMDAR encephalitis are linked to BOLD pattern incongruency. **(A)** Patients with NMDAR encephalitis showed reduced FC between the hippocampus and several DMN structures, including medial prefrontal and cingulate regions. While patients and controls showed similar levels of AUC-SCD in these edges, patients exhibited more incongruent BOLD signal patterns. **(B)** Correlation analyses between group-averaged functional measures. In hippocampal edges with reduced FC in patients, both HCs and patients have positive associations between FC and AUC-SCD and negative associations between FC and IPI. **(C)** Spearman’s rho correlation tests show that group-level alterations in FC are linked to complexity decreases (left) and IPI (right) across hippocampal connections.

Interestingly, in edges where patients showed reduced FC, we did not find significant differences in AUC-SCD compared to HC after correcting for multiple comparisons. Accordingly, patients with NMDAR encephalitis and HC exhibited similar levels of AUC-SCD in hippocampal edges, despite relatively strong reductions in hippocampal FC. Additionally, correlation analyses between group-average FC and AUC-SCD in these specific edges revealed even stronger associations in patients (ρ_L.hipp._ = 0.71, P_L.hipp._ = 0.013; ρ_R.hipp._ = 0.78, P_R.hipp._ < 0.001) than HCs (ρ_L.hipp._ = 0.52, P_L.hipp._ = 0.084; ρ_R.hipp._ = 0.62, P_R.hipp._ = 0.013). Given the strong positive relationship between FC and AUC-SCD seen both in HC and patients with NMDAR encephalitis across the whole brain (Figs. 2-3), we hypothesized that this finding may be explained by a diversification of the underlying BOLD signal patterns in patients.

Indeed, patients consistently showed increased IPI in the hippocampal edges with reduced FC compared to HC. This effect showed a slight lateralization towards the left hippocampus, where 8 of 12 edges with reduced FC showed significantly increased IPI in patients. In connections with the right hippocampus, the effect retained significance in 3 out of 16 edges after adjustment for multiple comparisons. Overall, connections with the most incongruent BOLD patterns involved the hippocampi bilaterally and the left and right thalamus, left cingulate gyrus, and right orbital gyrus (Table S2; Fig. 4a, right). We also performed correlation analyses between group-average FC and IPI in hippocampal edges with significant FC reductions. This revealed a strong inverse relationship between FC and IPI in both patients (ρ_L.hipp._ = −0.92, P_L.hipp._ < 0.0001; ρ_R.hipp._ = −0.95, P_R.hipp_. < 0.0001) and HCs (ρ_L.hipp._ = - 0.87, p_L.hipp._ < 0.001; ρ_R.hipp._ = −0.78, p_R.hipp._ < 0.001) (Fig. 4b). Moreover, we also observed a general relationship between lower FC in patients and lower AUC-SCD (left) as well as increased IPI (right) across all hippocampal edges – even in connections where between-group differences were limited to statistical tendency (Fig. 4c).

### Whole-brain FC reductions in NMDAR encephalitis are explained by BOLD pattern incongruency

While hippocampal FC reductions are a hallmark of NMDAR encephalitis, other FC changes have also been reported (*11*, *14*). Therefore, we extended the above approach to exploratory whole-brain (Fig. 5a) and network-level analyses (Fig. 5b). We observed significantly reduced FC that was predominantly localized to connections with deep grey matter structures. Specifically, connections of the thalamus were consistently affected in patients, with the strongest FC reductions between the thalamus and nucleus accumbens and orbital gyrus. In whole-brain analyses, patients exhibited several reductions in thalamic FC that were even stronger than those seen in analyses limited only to the hippocampus. These decreases in FC were paralleled by significant increases in IPI in patients, predominantly in connections between basal ganglia, cingulate gyri, and frontal regions of the DMN and FPN (Fig. 5a).

**Figure 5.**
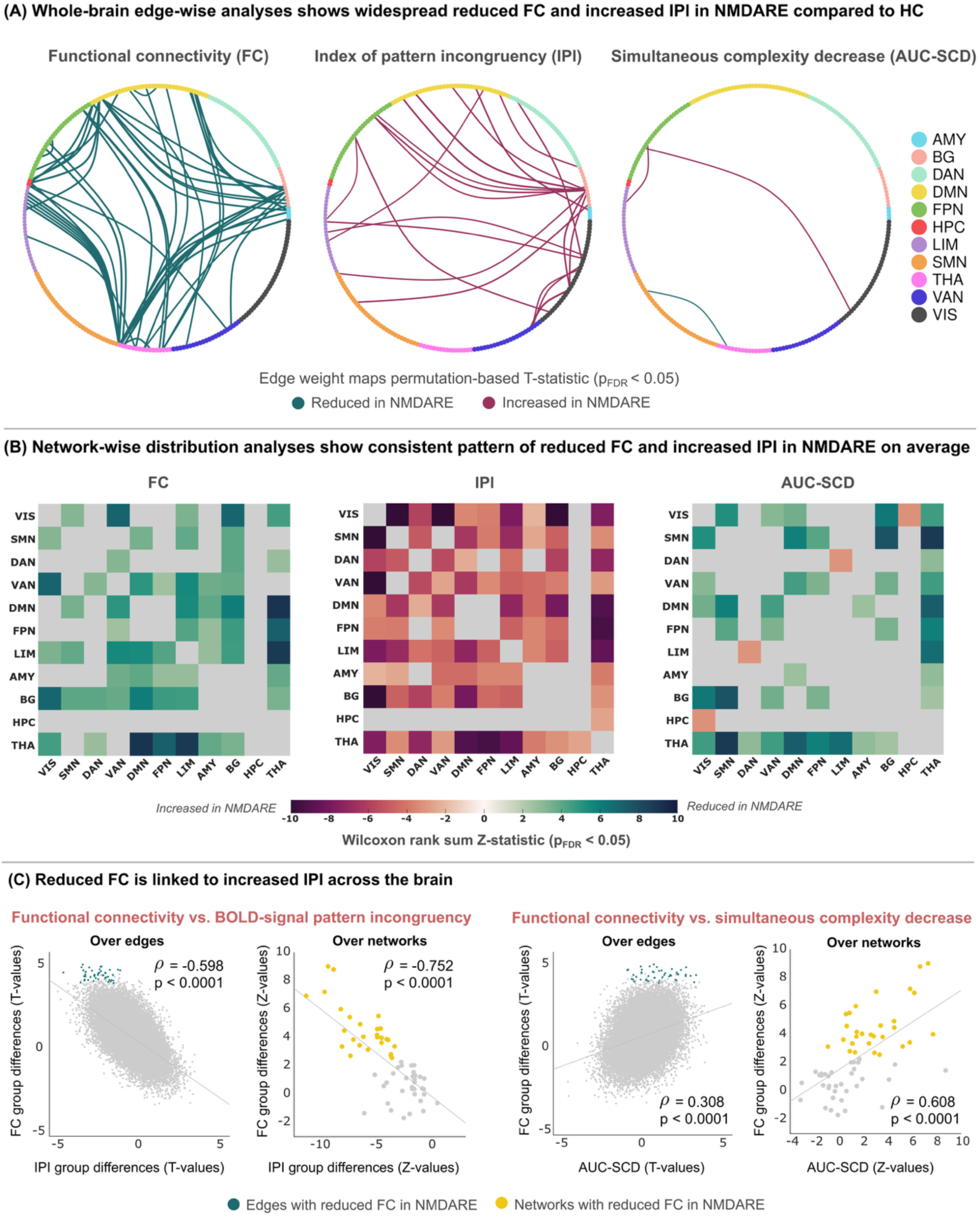
Whole-brain and network-specific FC reductions are linked to altered AUC-SCD and incongruent BOLD patterns. **(A)** Patients show widespread reductions in FC and increases in IPI. FDR-correction was computed over all edges. Nodes are color-coded according to RSN or subcortical region. **(B)** Network-wise analyses show a consistent pattern of reduced FC and increased IPI. Cells of the matrices map the Wilcoxon rank-sum Z-statistic. Greyed-out cells correspond to comparisons that did not survive FDR adjustment. **(C)** Correlations between group-level test statistics of FC changes and alterations of AUC-SCD and IPI, respectively.

To test the clustering of effects on the network level, we furthermore compared the network-averaged distributions of FC, IPI, and AUC-SCD between groups. In line with the above findings on hippocampal connectivity, patients consistently showed widespread reductions in FC and increases in IPI (Fig 5b). Specifically, the strongest FC reductions were found between the thalamus and the DMN (Z = 9.01, p < 0.0001), limbic network (Z = 8.80, p < 0.0001), and FPN (Z = 7.23, p < 0.0001), respectively. The strongest effects of increased IPI in patients were seen between the visual network and the VAN (Z = −12.39, p < 0.0001), the SMN (Z = −12.20, p < 0.0001), and the basal ganglia (Z = −10.94, p < 0.0001). IPI increases were also pronounced between thalamic connections with the FPN (Z = −9.33, p < 0.0001), DMN (Z = −9.01, p < 0.0001), and limbic network (Z = −8.53, p < 0.0001).

In terms of AUC-SCD, patients showed both increases and decreases compared to healthy participants. The strongest decreases were seen between the SMN and thalamus (Z = 8.76, p < 0.0001) and basal ganglia (Z = 7.72, p < 0.0001). Increases in AUC-SCD were found in connections between the visual network and hippocampus (Z = −3.16, p = 0.006) as well as between the limbic network and DAN (Z = −3.05, p = 0.008). Importantly, both whole-brain and network-level analyses confirmed the relationship between FC and complexity dynamics seen above in hippocampal analyses—lower FC was associated with higher IPI and lower AUC-SCD in patients, even when individual comparisons to HC were limited to statistical tendency (Fig. 5c).

### The normative relationship between AUC-SCD and IPI is disrupted in NMDAR encephalitis

Against the background of these FC findings, we next aimed to characterize the normative relationship between IPI and AUC-SCD. Specifically, we asked if this relationship is disrupted in patients and if such alterations can be localized to particular brain areas.

To this end, edgewise correlations were computed across HC to generate a spatial map of the relationship between IPI and AUC-SCD (Fig 6a, top). Across the brain, we observed predominantly inverse correlations between AUC-SCD and IPI, indicating that brain regions with more frequent or stronger simultaneous complexity decreases exhibit greater congruence in their BOLD signal patterns. In other words, when brain activity from two regions becomes more regular at the same time (i.e., a simultaneous complexity decrease), the symbolic patterns within those BOLD signals are more similar (i.e., low IPI). These inverse correlations were widespread across the brain, implicating connections from all networks and DGM regions. In contrast, we identified a much smaller subset of functional connections where IPI and AUC-SCD were *positively* correlated. Notably, a positive relationship in this context indicates that brain regions with more frequent or stronger simultaneous complexity decreases (i.e., high AUC-SCD), exhibit greater *incongruence* in their BOLD signal patterns (i.e., high IPI). Thus, when these regions exhibit complexity decreases at the same time, both underlying BOLD signals become more regular yet contain *different* distributions of symbolic patterns.

**Figure 6.**
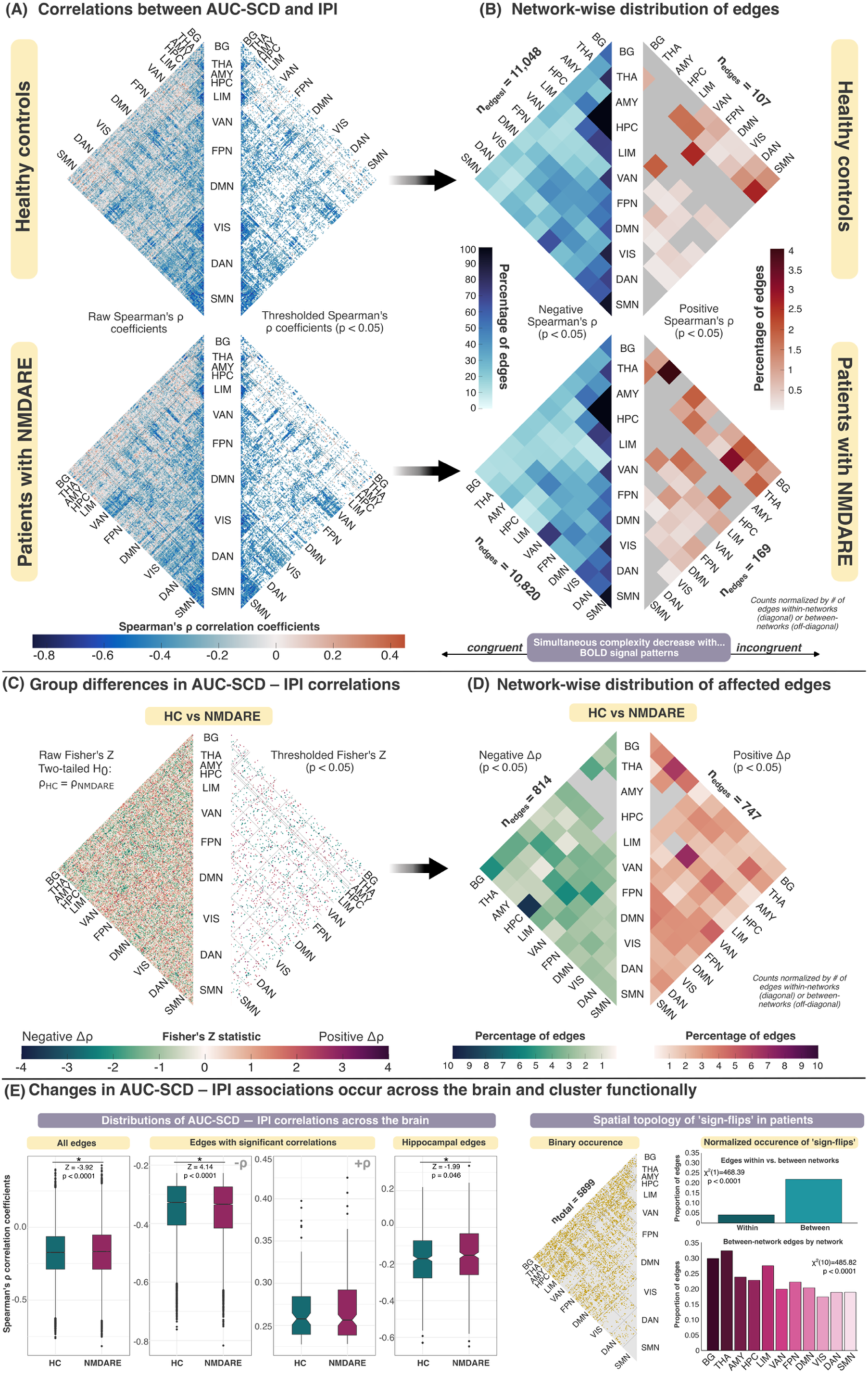
The normative relationship between IPI and AUC-SCD is disrupted in NMDAR encephalitis. **(A)** Whole-brain maps of IPI–AUC-SCD correlations in HCs (top) and patients (bottom). Left panels show all Spearman’s ρ coefficients and right panels show Spearman’s ρ coefficients significant at an alpha level of 0.05. Cells containing non-significant values are displayed in white. **(B)** Percentage of involvement in IPI–AUC-SCD effects, aggregated by network for HCs (top) and patients (bottom). Each cell of each triangle depicts the percentage of network involvement, calculated as the number of edges with negative (left) or positive (right) effects divided by the total number of corresponding edges. Only edges with significant IPI–AUC-SCD correlations were counted. **(C)** Whole-brain map of between-group differences in IPI–AUC-SCD correlations. Test statistics were generated for each edge using the Fisher’s Z-test for comparing correlations from independent groups, as implemented in the R ‘cocor’ package. Left panel shows all Z-statistics and right panel shows those thesholded at an alpha level of 0.05. **(D)** Percentage of involvement in IPI–AUC-SCD group differences, aggregated by network or DGM structure. As above, each cell depicts the percentage of network involvement, and edges with significant differences between groups were counted. **(E)** (Left) Distributions of IPI–AUC-SCD correlation coefficients over all edges, over edges with significant effects, and over hippocampal edges. Wilcoxon rank sum tests were used to compare between groups; (Right) Generalized disruption of normative effects summarized as ‘sign-flips’ (i.e., edges in which patients had an opposite direction of IPI–AUC-SCD correlation than HCs). Left matrix shows the binary occurrence of sign-flips across the brain; Right bar graphs show the proportion of edges where sign-flips occurred over all within-vs. between-network connections, as well as in RSN-specific connections.

Next, connections with significant correlations between IPI and AUC-SCD were aggregated by network (Fig 6b, top). Edges were categorized by the pairs of networks or DGM regions implicated in each, and the percentage of edges within and between each network that showed positive or negative IPI–AUC-SCD relationships was calculated. Visual inspection of network groupings revealed that within-network connections predominantly showed inverse correlations between IPI and AUC-SCD, while between-network connections showed a mixture of inverse and positive relationships. Edges with positive correlations accounted for up to 4% of all edges in any given network pair, while in some cases, inverse correlations were seen in 100% of edges. Interestingly, positive IPI–AUC-SCD correlations were primarily observed in connections from thalamic, BG, and limbic regions to networks involved in attention (VAN, DAN), cognition (FPN), and primary sensorimotor processing (SMN), while inverse correlations were seen across all networks and DGM regions.

Similarly to HCs, patients with NMDAR encephalitis showed predominantly inverse correlations between IPI and AUC-SCD across the brain, as well a subset of connections where IPI and AUC-SCD were instead positively related (Fig. 6a, bottom). However, patients showed a higher number of edges with significant positive correlations (n_PT_ = 169, n_HC_ = 107) and a lower number of edges with significant inverse correlations between IPI and AUC-SCD (n_PT_ = 10,820, n_HC_ = 11,048). On the network level, patients showed a higher percentage of thalamic, BG, and limbic edges with positive IPI–AUC-SCD correlations compared to HCs. Notably, hippocampal connections with DMN, FPN, and DAN regions showed positive IPI–AUC-SCD correlations in patients, while no hippocampal connections had positive correlations in HCs (Fig. 6b, bottom).

Between-group comparisons confirmed that patients with NMDAR encephalitis showed a widespread pattern of disrupted IPI–AUC-SCD associations, including both increases and decreases in correlation strength compared to HCs (Fig. 6c). When grouped by network, we observed that patient-specific alterations implicated nearly all pairs of networks and DGM structures (Fig. 6d). Furthermore, these disruptions were seen not only on the global scale (Z = −3.92, p < 0.0001), but also specifically across hippocampal connections, in which the distribution of IPI–AUC-SCD correlations was shifted positively in patients compared to HCs (Z = −1.99, p = 0.046; Fig. 6e, left).

Finally, we extended these analyses to consider the general case of edges in which patients showed an opposite effect direction in IPI–AUC-SCD relationships compared to HC. Given our aim to investigate patient-specific alterations, we furthermore studied the presence of ‘sign-flips’ in these associations relative to the HCs (Fig. 6e, right). Quantifying the binary occurrence of ‘sign-flips’ revealed that they tend to occur in connections with DGM structures and limbic regions, while cognitive and primary networks are less involved. For each network, we then computed the proportion of edges where sign-flips occurred in patients and used chi-square goodness-of-fit tests to evaluate if the observed occurrence of sign-flips deviated from the expected values, based on the proportional frequency of network labels in our population of edges. These analyses indeed confirmed that sign-flips occurred more frequently in between-network compared to within-network connections (X^2^(1) = 468.39, p < 0.0001), and that the occurrence of sign-flips is not proportionally distributed, but instead clusters by functional network (X^2^(10) = 485.82, p < 0.0001).

### BOLD pattern congruency in NMDAR encephalitis is associated with cognition beyond functional connectivity

Finally, we sought to explore how FC and our novel IPI measure respectively relate to cognitive performance in the patient cohort. To this end, we employed whole-brain correlation analyses to map associations between these brain measures and a composite cognition score across the domains of auditory-verbal memory, working memory, visual memory, executive functions, and attention (see Materials and Methods for details).

As expected, FC was positively related to cognitive performance, with strongest effects in limbic connections with the basal ganglia (Fig. 7a, left). In line with this, we consistently observed inverse relationships between IPI and cognition (Fig. 7a, right). Accordingly, better cognitive performance was associated with more congruent BOLD patterns, and this effect was present in several functional systems, including connections between regions of the limbic network and thalamus, default mode, basal ganglia and frontoparietal network.

**Figure 7.**
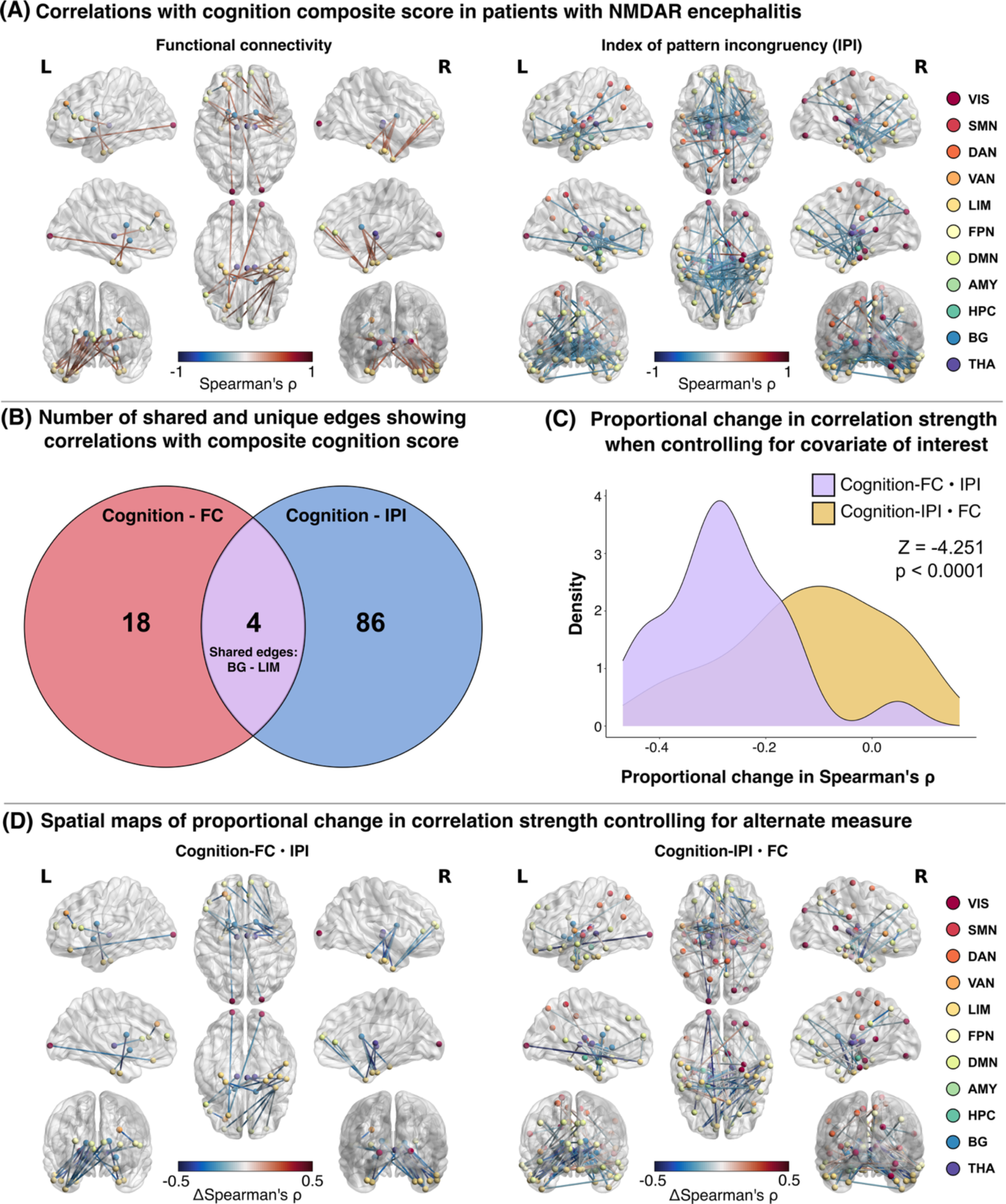
BOLD pattern congruency in NMDAR encephalitis is associated with cognition beyond functional connectivity. **(A)** Network visualizations of edges showing significant relationships between FC and cognition (left) and IPI and cognition (right). Visualizations were made with the BrainNet Viewer Toolbox for MATLAB. Edge color maps the strength and direction of Spearman’s rho correlation coefficients. Nodes are color-coded according to RSN or subcortical structure. **(B)** Venn diagram showing the number of unique and shared edges between the FC-cognition associations (left) and IPI-cognition associations (right). **(C)** Density plots showing the relative change in correlation strength when controlling for the respective other brain metric, tested with the Wilcoxon rank sum test. **(D)** Network visualizations depicting the spatial specificity of relative change in partial correlations. Edge color maps the direction and magnitude of the change in correlation strength between FC-cognition associations (left) and IPI-cognition associations (right) when controlling for the respective other brain measure.

Notably, comparisons between FC and IPI revealed that IPI maps additional correlations with cognition beyond those seen with FC (Fig. 7b). The intersection of both sets of associations contained four edges between the basal ganglia and limbic network. Additionally, we tested how the FC-cognition associations and the IPI-cognition associations would change when controlling for the respective other brain measure in partial correlations (i.e., FC-cognition when controlling for IPI and IPI-cognition when controlling for FC). We then calculated the percent change in correlation strength between the raw correlation coefficients and those calculated when controlling for the added covariate. We found that controlling for IPI reduced FC-cognition relationships to a significantly greater extent than controlling for FC in IPI-cognition relationships (Z = −4.25, p<0.001; Fig. 7c).

The strongest reductions were localized to basal ganglia and thalamic connections with the limbic network, where controlling for IPI reduced FC-cognition associations by 29-47%. Conversely, controlling for FC in IPI-cognition associations resulted in both increases and decreases in the strength of these associations. Increased effects after controlling for FC were not obviously clustered across the brain and ranged from 2-16%. Decreases due to controlling for FC-related variance were also widespread, with the strongest reductions seen in visual-DMN (46%) and basal ganglia-limbic connections (44%) (Fig. 7d).

## Discussion

In this study, we build upon the recently developed ‘complexome’ framework to show that functional connectivity (FC) is closely linked to the congruency of symbolic patterns in the underlying BOLD signals. To this end, we (i) integrate complexity dynamics with recent developments on signal cofluctuation analysis through edge-timeseries, (ii) extend the previous binary concept of simultaneous complexity drops to a continuous, threshold-free calculation (i.e., AUC-SCD), (iii) derive a new measure of pairwise dissimilarity in local BOLD signal patterns — the ‘index of pattern incongruency’ (IPI), (iv) show that the link between complexity dynamics and functional brain organization closely replicates in an independent dataset with different scanning parameters, and (v) provide first evidence of clinical relevance of this framework in the model disease of anti-N-methyl-D-Aspartate receptor (NMDAR) encephalitis. Specifically, we show that IPI captures unique information about complexity dynamics in addition to simultaneous complexity decreases (AUC-SCD), explains clinically relevant FC reductions in this patient group, and maps onto novel associations with cognitive performance beyond functional connectivity.

### Theoretical advances

The complexome framework was initially tested in the Human Connectome Project (HCP) Young Adults dataset, where we observed that resting-state brain activity is characterized by transient moments of high regularity in the BOLD signals, which become visible as spontaneous ‘complexity drops’. Importantly, we found that the FC strength between any two brain regions is closely related to how often they engage in these complexity drops at the same time (i.e., their ‘drop coincidence’). However, the definition of drop coincidence was based on a stringent binary drop threshold (the lower 1% of empirically observed complexity values across the sample). Consequently, a region was considered to show a complexity drop whenever the corresponding WPE value was low enough to meet this threshold. Similarly, two brain regions were considered to drop simultaneously if both signals independently reached the drop threshold in the same temporal window. While this binary formulation enabled many key insights into the spatial distribution, temporal propagation, and network effects of complexity drops (*1*), this approach necessitates an arbitrary drop threshold and is agnostic to the magnitude of the simultaneous complexity drops.

To overcome this binary definition, we here employed the recent mathematical framework of ‘edge-timeseries’ (*4*) and transferred this approach to cofluctuations between complexity timeseries. Instead of *BOLD signal* cofluctuations at the resolution of single timepoints, we thus obtained *complexity* cofluctuations representing neural dynamics at the resolution of temporal windows. To remain in line with the conceptualization of complexity *drops*, we then considered the instances of simultaneous *downward* complexity cofluctuations. This enabled us to capture both the duration (number of windows) and the magnitude (amplitude of cofluctuation) of simultaneous complexity decreases (SCD) in a single scalar as the area-under-the-curve (AUC-SCD) over time.

With this continuous SCD metric, we replicated a main finding from (*1*) in an independent dataset of healthy participants, confirming that AUC-SCD is strongly and positively related to FC strength. Importantly, this independent replication highlights that the relationship between FC and complexity dynamics is highly consistent across datasets, MRI machines, and scanning parameters.

Additionally, the extensions of the ‘complexome’ framework derived here allowed us to examine not only the coincidence of complexity decreases, but also the internal structure of the BOLD signals during those moments. In other words, by combining continuous measures of SCD and pairwise pattern incongruency, we were able to ask not just *when* regions become more regular together—but *how* their signal patterns relate in those moments.

To establish a normative account of these signal dynamics, we first focused on healthy participants. We observed that when activity from distinct regions becomes more regular, i.e. when two brains regions show complexity decreases at the same time, their BOLD signals show *similar* symbolic patterns. This inverse relationship (with IPI) was widespread, clustering especially *within* canonical RSNs and DGM structures. However, in about 1% of edges, we saw the opposite: regions decreasing in complexity at the same time displayed *different* underlying patterns. These connections were most common *between* DGM, limbic, and DMN regions to other cortical networks. Notably, however, a simultaneous complexity decrease with congruent or incongruent symbolic patterns represents the same phenomenon in an information-theoretical sense. Our findings therefore support a principled functional interpretation of the observed complexity dynamics, in line with our previous work (*1*). Simultaneous complexity decreases may express the informational architecture underpinning the brain’s functional network organization, whereas BOLD pattern incongruency may reflect the information content that is transmitted across this architecture. Our findings suggest that in healthy brain function these two aspects co-occur, while this relationship appears disrupted in neurological disease. Future work should investigate whether, and under what conditions, BOLD pattern incongruency is actively transmitted across the brain’s functional networks.

### Clinical application

The phenomenon of complexity drops has previously been shown to explain functional connectivity across the brain and was positively associated with cognitive and motor performance and negatively associated with age in healthy participants (*1*). Given this relationship, we sought to explain a well-established set of functional brain alterations in a clinical dataset. Patients with NMDAR encephalitis therefore served as a novel clinical application of our information-theoretic framework due to the highly replicable FC alterations seen in this population. Specifically, reduced FC between the hippocampus and the DMN is a well-established functional signature of NMDAR encephalitis (*9*, *11*, *14*) and was recently identified in murine models of the disease as well (*20*). Moreover, these functional changes are linked to reduced cognitive performance in NMDAR encephalitis, which persists for years after the acute stage of the disease (*9*, *21*). The localization of these alterations is thought to be linked to the especially high density of NMDA receptors in the hippocampal formation (*22*). In line with these previous reports, we here corroborated reduced hippocampal FC in NMDAR encephalitis, primarily in connections with the DMN and the thalamus.

Given the observed normative relationship between FC and AUC-SCD in the healthy brain, we hypothesized that these reductions in FC would be mirrored by parallel reductions in AUC-SCD. Interestingly, we found that patients and healthy participants showed similar levels of AUC-SCD among hippocampal connections. However, we previously found that the hippocampus was among those brain regions that engaged in complexity drops to a significantly lesser extent than other cortical areas (*1*). Therefore, we hypothesized that patients exhibit a shift in the symbolic patterns in the underlying BOLD signals that was not detected by complexity estimates alone. Indeed, we observed that hippocampal, whole-brain, and network-wise FC reductions in patients were systematically associated with increases in IPI. While some of these changes were also associated with AUC-SCD, effects tended to be even stronger for IPI. Relatedly, patients also showed widespread alterations in the association *between* IPI and AUC-SCD compared to healthy participants. While both groups had predominantly inverse correlations across the brain, patients with NMDAR encephalitis exhibited a positively skewed distribution of these correlations globally and had a higher percentage of edges with positive IPI–AUC-SCD correlations, especially in hippocampal and DGM connections. Accordingly, the normative IPI–AUC-SCD association seen in healthy individuals was disrupted in patients with NMDAR encephalitis, again underscoring the utility and clinical relevance of both measures. Besides these between-group effects, IPI also showed strong inverse associations with brain-wide FC within healthy participant and patient groups separately, suggesting a principled link between BOLD pattern congruency and BOLD signal covariance.

In addition, we also observed that IPI identified novel brain-cognition relationships in our clinical population. Specifically, more congruent BOLD patterns in basal ganglia and thalamic connections with the limbic and default mode networks were associated with better cognitive performance in patients. IPI-cognition associations were consistently inverse, i.e. higher similarity of symbolic patterns in BOLD signals was associated with better cognitive performance. IPI also mapped more associations with cognition compared to FC, underscoring its sensitivity to cognitive performance. Furthermore, controlling for IPI in FC-cognition relationships significantly reduced the strength of these associations, while the inverse did not equivalently hold.

Overall, these findings clearly support the view that NMDAR encephalitis is characterized by increased volatility of functional brain dynamics (*12*, *13*), and that this volatility is rooted in altered complexity dynamics.

### Future directions

Here, we limited FC analyses to static accounts of BOLD activity. However, the new continuous definition of SCD will facilitate future research into the moment-to-moment relationships between FC and complexity dynamics. Moreover, our study focused on group-level effects, but our framework naturally extends to the study of intra-individual differences. This is especially interesting in the context of precision neuroimaging datasets to explore the variability of complexity dynamics over days to weeks.

Furthermore, the normative analyses in our study carry several immediate implications. One the one hand, future studies in lifespan datasets will be able to derive a normative account of complexity dynamics in human development. On the other hand, the framework is easily applicable to other clinical conditions as well, lending the opportunity to investigate disease-specific alterations in functional brain dynamics. The demonstrated sensitivity of IPI to elucidate associations between brain dynamics and cognitive symptoms that otherwise remain undetected suggests that this measure may be useful for investigations of other neuropsychiatric disorders as well. Future applications to other patient populations, for example those with complex neuropsychiatric symptoms (i.e., schizophrenia) or other autoimmune-mediated conditions (i.e., multiple sclerosis, LGI-1 encephalitis), would be especially interesting to evaluate the disease-specificity of the present findings.

Overall, the present study provides an in-depth replication, theoretical extension, and one of the first clinical applications of the recently developed ‘complexome’ framework. We introduce a novel measure of BOLD pattern incongruency that is closely linked to FC and cognitive performance, and demonstrate that both complexity cofluctuations and BOLD pattern incongruency provide unique yet complementary insights into the dynamics of human brain activity. Taken together, these findings underscore the value of IPI as a novel and informative measure of BOLD signal dynamics, with relevance not only for advancing our normative understanding of brain organization, but also for identifying deviations in clinical populations.

## Materials and Methods

### Participants and clinical assessment

We studied a sample of 75 patients with anti-NMDA receptor (NMDAR) encephalitis and 75 age- and sex-matched healthy control participants (HCs). Patients were recruited from the Department of Neurology at Charité-Universitätsmedizin Berlin. Healthy control participants (HCs) were recruited using the Charité Intranet. Diagnosis of NMDAR encephalitis was based on clinical presentation and the detection of immunoglobulin G (IgG) NMDA receptor antibodies. Patients were in the post-acute stage of the disease with an average time since disease onset of 2.97 ± 2.48 years. The study was approved by the local ethics committee [EA1/206/10, EA1/095/12, EA1/105/16, EA4/011/19], and all participants gave written informed consent.

For cognitive assessment, patients underwent a battery of standardized neuropsychological tests. A composite cognition score was calculated using data from patients with complete cognitive neuropsychological test scores. Out of 75 patients, 61 had complete data and were included in final brain-cognition correlations. We selected two scores from each of the following cognitive sub-domains: auditory-verbal memory (Rey Auditory Verbal Learning Test summary and delayed recall), working memory (Digit Span Forward and Digit Span Backward), visual memory (Rey-Oesterreich Complex Figure immediate and delayed-recall), executive function (Stroop test, Go-No-Go test), and attention (Test of Attentional Performance divided attention auditory and divided attention visual). Scores that corresponded to reaction times were re-coded by multiplying by –1 to maintain the directionality of the composite score, such that higher scores represent better cognitive performance relative to the mean across all patients. For each of the 10 tests, a z-score was calculated over the patient scores. The mean was then taken over a patients’ 10 z-scores, resulting in a mean cognitive performance composite z-score for each patient.

### MRI data acquisition

MRI data were acquired at the Berlin Center for Advanced Neuroimaging at Charité-Universitätsmedizin Berlin, Germany, using a 20-channel head coil on a 3 T Trim Trio scanner (Siemens, Erlangen, Germany). For each participant, the following sequences were acquired for this study: a 10-minute resting-state-fMRI (rsfMRI) scan was collected using a repetition time (TR) of 2250 ms (TE = 30 ms, 260 volumes, matrix size = 64 × 64, 37 axial slices, slice thickness = 3.4 mm, voxel size = 3.4 × 3.4 × 3.4 mm^3^), and a T1-weighted structural scan was acquired using a magnetization-prepared rapid gradient echo (MPRAGE) sequence (1 mm^3^ isotopic resolution, matrix size = 240 × 240, 176 slices).

### Resting-state fMRI preprocessing

Prior to preprocessing, framewise displacement (FD) was calculated for each participant and assessed against a mean FD cutoff of 0.50mm (*23*). No participants had a mean FD greater than or equal to 0.50mm. Preprocessing was performed using customized MATLAB scripts based on Parkes et al. (*24*) and included removal of the first 4 volumes of each participant’s rs-fMRI scan, slice-timing correction, detrending of BOLD timeseries, intensity normalization, and spatial smoothing by convolving with a 6mm full width at half maximum Gaussian kernel. Motion correction was performed by regression of six realignment parameters, mean white matter and CSF signals. Bandpass filtering to retain frequencies between 0.01 and 0.1 Hz and demeaning were performed. The Human Brainnetome Atlas (*16*) was customized to combine lateral hippocampal regions of interest (ROIs) into one left and one right hippocampal ROI, resulting in 210 cortical and 34 subcortical ROIs in total. Timeseries were volumetrically extracted by averaging across BOLD signals from every voxel corresponding to a given ROI.

### Weighted permutation entropy calculation

Time-resolved WPE was calculated using custom MATLAB scripts as described in detail in (*1*). Code for the calculation of WPE was adapted from the online repository originally published with (*1*) (https://osf.io/mr8f7/). WPE is an information-theoretic measure of signal complexity based on a symbolic-encoding framework (*5*). Therein, the regularity of a timeseries is estimated through the occurrence of possible abstract patterns over time (i.e., the pattern frequency distribution) (*1*, *5*). The mathematical formula for WPE is:

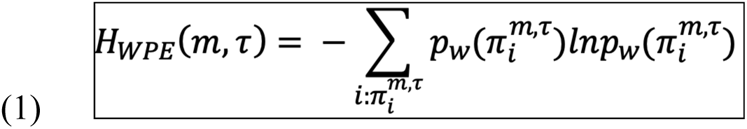

To remain in line with recent applications of WPE on neural data (*1*), we selected a motif length of m = 3 and a lag parameter of tau = 1. The motif length, m, is the number of consecutive time points in a signal that are simultaneously transformed into rank space by evaluating the position of each data point in relation to the m-1 other timepoints. The motif length directly determines the number of possible patterns present in the timeseries. For example, a motif length of m = 3 results in 6 (i.e., factorial of 3) possible patterns: [1,2,3], [1,3,2], [2,1,3], [2,3,1], [3,1,2], [3,2,1]. The lag parameter τ is the number of time points by which each consecutive rank encoding step progresses forward along the timeseries. Over a given window, the rank transformation, motif encoding, and progression steps are iteratively performed along the timeseries, resulting in a frequency distribution across the possible patterns. Importantly, amplitude information is incorporated into frequency distributions by a weighting factor that expresses the proportion of amplitude variance accounted for by each motif (*5*). This amplitude-weighted pattern frequency distribution is then normalized to produce the weighted relative frequencies for each motif which sum to unity. Finally, WPE is calculated as the Shannon entropy over this distribution and qualified by log_2_(m!), producing normalized entropy values between 0 and 1.

Here, we implemented a time-resolved calculation of WPE using a sliding window technique, which has been described in detail in (*1*) and is commonly used in the dynamic FC literature (*24*, *25*). Briefly, each BOLD timeseries was segmented into overlapping windows with a window size of 20 TR (45 seconds) and a slide length of 1 TR (2.25 seconds). The application of this technique resulted in a set of 233 windows (95% overlap) for each BOLD timeseries. For every ROI and within every window, WPE was calculated on the 20 second timeseries snippet. This resulted in a WPE timeseries (i.e., complexity timeseries) for every ROI with 233 data-points at the resolution of windows. These parameters mirrored the window length as applied in the HCP data (*1*), accounting for significant differences in the temporal resolution of our in-house fMRI data (TR: 2.25s) compared to that of the HCP (TR: 0.72s). Despite these differences in temporal resolution, complexity estimates in our in-house cohort of healthy participants were comparable to those in the HCP dataset, and relationships between complexity dynamics and FC were remarkably consistent.

### Complexity timeseries cofluctuation

Key insights of the original complexome framework included the following findings: (1) the brain tends to operate in a state of high complexity, which is seen ubiquitously across the cortex and subcortex; (2) brain regions spontaneously engage in complexity drops, which represent transient moments of increased regularity in the BOLD signals; (3) the presence of complexity drops across the brain is not random, but follows a spatially and temporally structured architecture. Subcortical regions engage in complexity drops less frequently than cortical areas, and drop affinity within the cortex follows a unimodal-to-transmodal gradient (*1*).

To characterize the spatiotemporal pattern of complexity dynamics across the brain, we implemented a pairwise measure referred to as “drop coincidence”. This measure quantifies the number of instances (windows) in which a pair of brain regions simultaneously engage in a complexity drop. Therein, the definition of a complexity drop was determined based on the lowest 1% threshold of WPE values across all empirically observed values, such that any complexity value meeting this threshold in a given window was considered a complexity drop. While this implementation of drop coincidence yielded significant insight into the spatiotemporal complexity architecture of brain activity, several modifications are useful to reduce the reliance on a priori defined thresholds and binarization of drop coincidence between brain regions.

Thus, in the present work, we developed a novel computation that enables a continuous account of simultaneous complexity changes. To this end, we employed the recent edge-timeseries framework from (*4*) to compute a pairwise measure of complexity timeseries *cofluctuations* at the resolution of temporal windows. The edge-timeseries framework represents a new method that decomposes functional connectivity into frame-wise contributions to the overall timeseries correlation.

Here, we transferred this approach to the realm of complexity dynamics: For every pair of complexity timeseries, we performed the following steps to compute an analogous measure of ‘complexity cofluctuation’:

a. Each complexity timeseries was z-scored. The element-wise product between a pair of z-scored complexity timeseries was taken as the edge-timeseries (i.e., complexity cofluctuation timeseries).
b. The root sum square of this edge-timeseries was calculated and taken as the amplitude of complexity cofluctuation at the resolution of temporal windows.

Within the above framework, a positive value in the edge-timeseries represents a window in which the underlying complexity timeseries fluctuate in the same direction (i.e., both move upward *or* downward). Given the previously established definition of complexity drops, we then filtered the edge-timeseries to select only the instances corresponding to simultaneous *decreases* in complexity (SCD). An instantaneous decrease in complexity was defined within a complexity timeseries, such that in window *t* the complexity value was less than in window *t-1*. Using these indices of downward cofluctuations, we then extracted the corresponding amplitude of the cofluctuation timeseries. The area-under-the-curve (AUC) of the thus-indexed timeseries was then taken as a summary measure over time, incorporating both the magnitude (amplitude of cofluctuation) and duration (number of temporal windows) of SCD between any pair of brain regions.

### Index of pattern incongruency

To quantify the differences in symbolic patterns between a pair of BOLD signals, we derived a novel measure called the ‘index of pattern incongruency’ (IPI). For each instance of SCD, we computed the Euclidean distance between the pattern frequency distributions of the two corresponding BOLD windows, as obtained during the calculation of WPE. The Euclidean distance between pairs of probability distributions was implemented according to (*26*) (see https://github.com/preethamam/pdfsDistanceandSimilarity). This results in a scalar distance value that captures the dissimilarity between the symbolic patterns present in the two BOLD signals. Finally, the mean over all indexed windows was taken as the IPI between a given pair of brain regions.

Given the mathematical formulation of Euclidean distance, the analytical bounds of IPI are 0 and ✓2. An IPI of 0 indicates that, in every window of simultaneous downward complexity cofluctuations, a pair of BOLD signals have identical pattern frequency distributions. Accordingly, an IPI of ✓2 indicates that, in *every* window of simultaneous downward complexity cofluctuation, a pair of BOLD signals have maximally distinct pattern frequency distributions. In the case of normalized probability distributions (i.e., each summing to unity), the Euclidean distance between n-dimensional distributions approaches the upper bound of ✓2 as the maximum value present in the distribution approaches 1. Herein, a Euclidean distance of exactly ✓2 is theoretically possible only when, within a given window, each BOLD signal’s pattern is explained entirely by one motif *and* that motif is different for each signal. Thus, an IPI of exactly ✓2 is theoretically possible only when, in *every* window of simultaneous downward complexity cofluctuation, a pair of BOLD signals each have deterministic, *yet* distinct pattern frequency distributions. Empirically, we observe a somewhat narrower range of window-wise Euclidean distances during moments of SCD [0.00041, 1.35], at least partially due to the rarity of BOLD signals exhibiting deterministic behavior; WPE = 0 in less than 0.00005% of instances. Since IPI also accounts for the number of windows in which SCD occurs, we again observed slightly more narrow empirical bounds [0.031, 0.65].

Taken together, this novel IPI measure serves as an extension of time-resolved BOLD signal complexity analysis with WPE. Since the WPE algorithm is based on the Shannon entropy of the empirical pattern distribution, it is agnostic to which particular pattern drives the signal. Therefore, it is possible for two different pattern distributions to yield equal WPE values – because, in an information-theoretic sense, they are the same (e.g., consider a completely deterministic distribution that accumulates the whole probability mass in pattern [1 2 3] and another one with pattern [2 3 1], which both yield an entropy of zero). In contrast, our definition of the Euclidean distance between two distributions is sensitive to differences in the frequency of each possible pattern present in the signal. For an extended explanation, see Supplementary Text 1.

### Static functional connectivity

The product-moment correlation coefficient was computed between every pair of ROI timeseries to create a 244-by-244 static functional connectivity (FC) matrix for each participant. Raw correlation coefficients were Fisher r-to-z-transformed by taking the element-wise inverse hyperbolic tangent of each participants’ FC matrix.

### Statistical Analysis

#### FC–IPI and FC–AUC-SCD analyses

Within-group correlation analyses were calculated with Spearman’s rank tests on mean FC, IPI, and AUC-SCD. Multiple regression and commonality analyses were employed to evaluate the normative relationship between FC, IPI, and AUC-SCD, using the “yhat” package for R (*19*). Commonality analysis is a method for partitioning the variance explained in a dependent variable by multiple independent variables into unique and shared portions. Herein, commonality analysis was used to quantify the variance in FC that is explained by IPI and AUC-SCD in healthy participants.

Group-level differences in whole-brain and hippocampal FC, IPI, and AUC-SCD were computed using a non-parametric, permutation-based T-test (see https://version.aalto.fi/gitlab/BML/bramila). Group differences in network-wise brain measures were computed on the group averages to maintain spatial specificity. For each pair of networks, the mean values for a given edge were computed, and Wilcoxon rank sum tests over network edges were used to test for differences between HC and NMDAR encephalitis. In case of multiple univariate comparisons, the false discovery rate (FDR) was controlled using the Benjamini-Hochberg procedure (*27*).

#### IPI–AUC-SCD analyses

Edgewise Spearman’s rank correlation tests between measures of IPI and AUC-SCD were performed for each group, calculating a coefficient over the IPI and AUC-SCD values for each edge, resulting in a whole-brain correlation map for each group. Whole-brain maps were then thresholded, and edges with significant IPI–AUC-SCD correlations underwent network-wise aggregation to summarize the extent to which each network was affected. Network-wise percentages were calculated by counting the number of edges that had a positive or negative correlation and dividing by the total number of edges within or between a network/region. This resulted in a matrix of percentages, where values on the diagonal represent the percentage of edges within each network that met a given criteria and values off the diagonal represent percentages between pairs of networks.

Edgewise between-group comparisons of IPI–AUC-SCD correlations were performed using the R ‘cocor’ package (*28*). Specifically, we applied Fisher’s z-test for comparing two correlation coefficients from independent groups (*29*). This entails standardizing the correlation coefficients using Fisher’s r-to-Z transformation (*30*) and then calculating the z-statistic given the sample size of each group. According to (*31*), the application of Fisher’s r-to-Z transformation on Spearman’s ρ rather than Pearson’s r coefficients is reliable, despite non-parametric assumptions. Between-group comparisons of distributions of correlation coefficients were performed using Wilcoxon rank sum tests.

Finally, to analyze ‘sign-flips’ of IPI–AUC-SCD associations in patients, we computed the normalized proportion of edges where patients showed opposite effect directions compared to HC by summing the number of occurrences and dividing by the total number of edges. This was calculated independently for all within- and between-network connections, as well as for each network and DGM region. We performed chi-square goodness-of-fit tests to evaluate if the observed occurrence of sign-flips deviated from the expected values, based on the proportional frequency of network and structure labels in our population of edges. Due to the exploratory nature of these analyses in particular, statistical results are reported at an uncorrected alpha level of 0.05.

#### Brain-cognition analyses

Correlation analyses between cognition scores and brain measures were performed using data from patients with complete neuropsychological data (n = 61/75). All correlations were computed using Spearman’s rank correlation tests and, where applicable, Spearman’s rank partial correlation tests. The proportion of change in correlation strength between full and partial correlations was calculated for each edge. This was computed as the absolute value of the difference between a given partial ρ and its respective full ρ, divided by the absolute value of the full ρ. The resultant was then multiplied by the sign of the absolute value of the partial ρ minus the absolute value of the full ρ. Thus, the proportional change in correlation strength captures both the magnitude of change with respect to the strength of correlation, as well as the directionality of both underlying coefficients. Due to the exploratory nature of this approach, we thresholded correlation tests at a univariate α-level of 0.001.

## Acknowledgements

None.

## Author contributions

Conceptualization: AR, NvS, FP, HP, SK, CF

Data curation: AR, NvS

Formal analysis: AR

Funding acquisition: CF

Investigation: AR, SK

Methodology: AR, NvS, SK

Resources: FP, HP, CF

Software: AR, SK

Supervision: SK, CF

Validation: AR, SK

Visualization: AR

Writing—original draft: AR, SK, CF

Writing—review and editing: AR, NvS, FP, HP, SK, CF

## Data and materials availability

Anonymized brain derivatives and analysis code to reproduce the findings of this study will be made available on the Open Science Framework. Upon reasonable request, clinical data can be provided pending scientific review and a completed material transfer agreement. Requests should be submitted to.

## References

1. S. Krohn, N. von Schwanenflug, L. Waschke, A. Romanello, M. Gell, D. D. Garrett, C. Finke, A spatiotemporal complexity architecture of human brain activity. Science Advances 9 (2023).

2. L. Waschke, N. A. Kloosterman, J. Obleser, D. D. Garrett, Behavior needs neural variability. Neuron 109, 751–766 (2021).

3. B. Biswal, F. Zerrin Yetkin, V. M. Haughton, J. S. Hyde, Functional connectivity in the motor cortex of resting human brain using echo-planar mri. Magn. Reson. Med. 34, 537–541 (1995).

4. F. Z. Esfahlani, Y. Jo, J. Faskowitz, L. Byrge, D. P. Kennedy, O. Sporns, R. F. Betzel, High-amplitude cofluctuations in cortical activity drive functional connectivity. PNAS 117, 28393–28401 (2020).

5. B. Fadlallah, B. Chen, A. Keil, J. Príncipe, Weighted-permutation entropy: A complexity measure for time series incorporating amplitude information. Phys. Rev. E 87, 022911 (2013).

6. W. W. Seeley, R. K. Crawford, J. Zhou, B. L. Miller, M. D. Greicius, Neurodegenerative Diseases Target Large-Scale Human Brain Networks. Neuron 62, 42–52 (2009).

7. M. M. Schoonheim, T. A. A. Broeders, J. J. G. Geurts, The network collapse in multiple sclerosis: An overview of novel concepts to address disease dynamics. NeuroImage: Clinical 35, 103108 (2022).

8. M. Peer, H. Prüss, I. Ben-Dayan, F. Paul, S. Arzy, C. Finke, Functional connectivity of large-scale brain networks in patients with anti-NMDA receptor encephalitis: an observational study. The Lancet Psychiatry 4, 768–774 (2017).

9. C. Finke, U. A. Kopp, M. Scheel, L.-M. Pech, C. Soemmer, J. Schlichting, F. Leypoldt, A. U. Brandt, J. Wuerfel, C. Probst, C. J. Ploner, H. Prüss, F. Paul, Functional and structural brain changes in anti-N-methyl-D-aspartate receptor encephalitis. Ann Neurol. 74(2), 284–96 (2013).

10. J. Dalmau, E. Lancaster, E. Martinez-Hernandez, M. R. Rosenfeld, R. Balice-Gordon, Clinical experience and laboratory investigations in patients with anti-NMDAR encephalitis. The Lancet Neurology 10, 63–74 (2011).

11. N. von Schwanenflug, S. Krohn, J. Heine, F. Paul, H. Prüss, C. Finke, State-dependent signatures of anti-N-methyl-D-aspartate receptor encephalitis. Brain Communications 4 (2022).

12. N. von Schwanenflug, J. P. Ramirez-Mahaluf, S. Krohn, A. Romanello, J. Heine, H. Prüss, N. A. Crossley, C. Finke, Reduced resilience of functional state transitions in patients with anti-NMDA receptor encephalitis. doi: 10.1101/2022.01.24.477081 (2022).

13. T. J. Hartung, N. von Schwanenflug, S. Krohn, T. A. A. Broeders, H. Prüss, M. M. Schoonheim, C. Finke, Eigenvector centrality mapping reveals volatility of functional brain dynamics in anti-NMDA receptor encephalitis. Biological Psychiatry: Cognitive Neuroscience and Neuroimaging, doi: 10.1016/j.bpsc.2024.07.021 (2024).

14. M. Peer, H. Prüss, I. Ben-Dayan, F. Paul, S. Arzy, C. Finke, Functional connectivity of large-scale brain networks in patients with anti-NMDA receptor encephalitis: an observational study. The Lancet Psychiatry 4, 768–774 (2017).

15. L. Cai, Y. Liang, H. Huang, X. Zhou, J. Zheng, Cerebral functional activity and connectivity changes in anti-N-methyl-D-aspartate receptor encephalitis: A resting-state fMRI study. NeuroImage: Clinical 25, 102189 (2020).

16. L. Fan, H. Li, J. Zhuo, Y. Zhang, J. Wang, L. Chen, Z. Yang, C. Chu, S. Xie, A. R. Laird, P. T. Fox, S. B. Eickhoff, C. Yu, T. Jiang, The Human Brainnetome Atlas: A New Brain Atlas Based on Connectional Architecture. Cereb. Cortex 26, 3508–3526 (2016).

17. D. C. Van Essen, K. Ugurbil, E. Auerbach, D. Barch, T. E. J. Behrens, R. Bucholz, A. Chang, L. Chen, M. Corbetta, S. W. Curtiss, S. Della Penna, D. Feinberg, M. F. Glasser, N. Harel, A. C. Heath, L. Larson-Prior, D. Marcus, G. Michalareas, S. Moeller, R. Oostenveld, S. E. Petersen, F. Prior, B. L. Schlaggar, S. M. Smith, A. Z. Snyder, J. Xu, E. Yacoub, The Human Connectome Project: A data acquisition perspective. Neuroimage 62, 2222–2231 (2012).

18. T. B. T. Yeo, F. M. Krienen, J. Sepulcre, M. R. Sabuncu, D. Lashkari, M. Hollinshead, J. L. Roffman, J. W. Smoller, L. Zöllei, J. R. Polimeni, B. Fischl, H. Liu, R. L. Buckner, The organization of the human cerebral cortex estimated by intrinsic functional connectivity. J Neurophysiol 106, 1125–1165 (2011).

19. K. Nimon, M. Lewis, R. Kane, R. M. Haynes, An R package to compute commonality coefficients in the multiple regression case: An introduction to the package and a practical example. Behavior Research Methods 40, 457–466 (2008).

20. J. Kuchling, B. Jurek, M. Kents, J. Kreye, C. Geis, J. Wickel, S. Mueller, S. P. Koch, P. Boehm-Sturm, H. Prüss, C. Finke, Impaired functional connectivity of the hippocampus in translational murine models of NMDA-receptor antibody associated neuropsychiatric pathology. Mol Psychiatry, doi: 10.1038/s41380-023-02303-9 (2023).

21. J. Heine, U. A. Kopp, J. Klag, C. J. Ploner, H. Prüss, C. Finke, Long-Term Cognitive Outcome in Anti–N-Methyl-D-Aspartate Receptor Encephalitis. Annals of Neurology 90, 949–961 (2021).

22. D. Monaghan, C. Cotman, Distribution of N-methyl-D-aspartate-sensitive L-[3H]glutamate-binding sites in rat brain. J. Neurosci. 5, 2909–2919 (1985).

23. J. Power, K. Barnes, A. Snyder, B. Schlaggar, S. Petersen, Spurious but systematic correlations in functional connectivity MRI networks arise from subject motion. NeuroImage 59, 2142– 2154 (2012).

24. L. Parkes, B. Fulcher, M. Yücel, A. Fornito, An evaluation of the efficacy, reliability, and sensitivity of motion correction strategies for resting-state functional MRI. NeuroImage 171, 415–436 (2018).

25. E. A. Allen, E. Damaraju, S. M. Plis, E. B. Erhardt, T. Eichele, V. D. Calhoun, Tracking Whole-Brain Connectivity Dynamics in the Resting State. Cerebral Cortex 24, 663–676 (2014).

26. S.-H. Cha, Comprehensive Survey on Distance/Similarity Measures between Probability Density Functions. International Journal of Mathematical Models and Methods in Applied Sciences 1 (2007).

27. Y. Benjamini, Y. Hochberg, Controlling the False Discovery Rate: A Practical and Powerful Approach to Multiple Testing. Journal of the Royal Statistical Society. Series B (Methodological) 57, 289–300 (1995).

28. B. Diedenhofen, J. Musch, cocor: A Comprehensive Solution for the Statistical Comparison of Correlations. PLoS ONE 10, e0121945 (2015).

29. R. A. Fisher, Statistical Methods for Research Workers (Oliver and Boyd, Edinburgh, Scotland, 1925; https://psychclassics.yorku.ca/Fisher/Methods/).

30. R. A. Fisher, On the probable error of a coefficient of correlation deduced from a small sample. Metron 1, 1–32 (1921).

31. L. Myers, M. J. Sirois, Spearman Correlation coefficients, differences between, Encyclopedia of Statistical Sciences (2006). 10.1002/0471667196.ess5050.pub2.

