## Supplementary Materials for "Functional connectivity is linked to symbolic BOLD patterns: replication, extension, and clinical application of the human ‘complexome’"

Amy Romanello *et al.*

Corresponding author: Carsten Finke.


**Supplementary Text**

A note on weighted permutation entropy and the occurrence of individual motifs

The mathematical formulation of weighted permutation entropy (WPE) relies on transforming the individual datapoints within a BOLD timeseries into rank space. We here employed a motif length of m = 3. In other words, three consecutive datapoints are considered and categorized based on their rank structure into one of 6 (m factorial) possible permutations. This categorization step (i.e., rank-encoding and amplitude weighting) is repeated along the timeseries, sliding by 1 datapoint and considering 3 datapoints each time, until the number of occurrences of each possible motif is counted. This results in a probability distribution that depicts the frequency of each pattern that occurred in the BOLD signal during a given timeframe. The Shannon entropy is then computed on this distribution, with the resulting value serving as the WPE estimation.

Within a given signal, this framework quantifies the amount of information encoded in the signal. We here apply a *time-resolved* calculation of WPE, using a sliding window method to segment each BOLD signal into overlapping windows. Thus, *within a given window*, a BOLD signal that is constantly increasing as it approaches an inflection point will have a high frequency of the [1, 2, 3] motif and low frequencies for every other pattern, whereas another BOLD signal that is constantly decreasing after an inflection point will have a high frequency of the [3, 2, 1] motif and low frequencies for every other pattern. Inflection points in the signal are characterized by the rarer motifs: [1, 3, 2], [2, 3, 1], [2, 1, 3], and [3, 1, 2] (1).

In this toy example, let the frequency distributions of these two signal snippets (monotonically increasing, monotonically decreasing) be exact mirror opposites of one another. Importantly, in the information theoretic sense, both exemplar signals are considered to be highly predictable and carry a low amount of information; in fact, their WPE values are close to zero and exactly the same. The WPE algorithm is not sensitive to the occurrence of individual motifs within a pattern frequency distribution, and thus quantifies the amount of information carried over the distribution of all possible patterns. On the contrary, the toy BOLD signals described above would necessarily have a low level of covariance and thus a FC estimate close to zero. This exemplar scenario presents a case in which the synchronization between a pair of BOLD signals is low, yet in the information theoretic sense, the signals are equally complex (or *not* complex). This toy example underscores the utility of a novel metric that captures information about the similarity of BOLD patterns and that which can be used to link underlying signal properties to both changes in nodal complexity and connectivity between signals. Our so-called ‘index of pattern incongruency’ (IPI) aims to address this gap.

### Fig. S1.

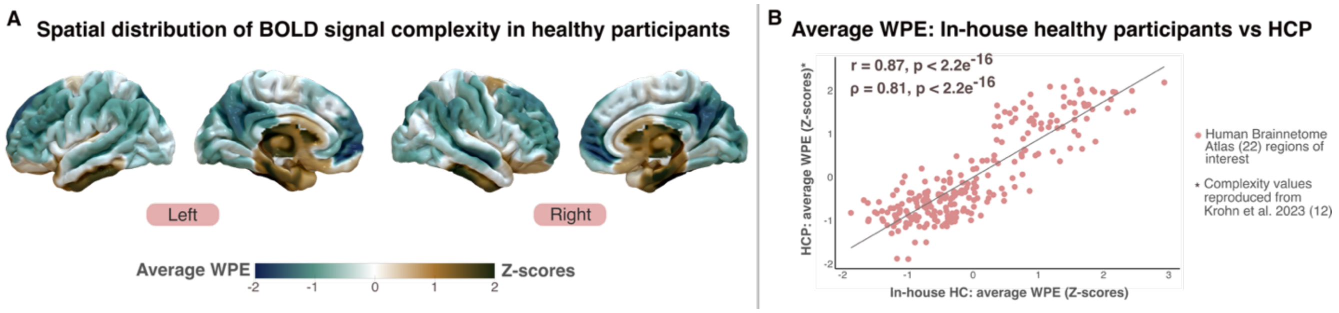


**Figure S1. Average signal complexity of in-house healthy participants is consistent with findings from the HCP Young Adult dataset. (A)** Mean BOLD signal complexity values (WPE) computed over healthy control participants (n = 75) and mapped to brain space. Mean WPE values were Z-scored over regions. Gold colors represent high complexity and blue colors represent low complexity. **(B)** Correlation between the average complexity values displayed in (A) and the re-computed average complexity values from the Human Connectome Project (HCP) dataset, originally analyzed in (1). Both analyses applied the same parcellation schema using the Human Brainnetome Atlas, thus the spatial topologies of WPE values could be directly compared. Strong positive correlation coefficients indicate that the spatial distribution of WPE is highly consistent between datasets.

**Table S1. Hippocampal functional connectivity alterations in patients with NMDAR encephalitis.**

| Seed region | Anatomical label | Detailed anatomical and modified cyto-architectonic description | Network label | MNI coordinates (X, Y, Z) | T | $p_{uncor}$ | $p_{FDR}$ | Effect size (Cohen's D) |
|---|---|---|---|---|---|---|---|---|
| L. HPC | L. sup. frontal gyrus | A9m, medial area 9 | DMN | -5, 36, 38 | 3.12 | <0.001 | 0.026 | 0.399 |
| | L. sup. frontal gyrus | A10m, medial area 10 | DMN | -8, 56, 15 | 3.74 | <0.001 | 0.004 | 0.459 |
| | R. sup. frontal gyrus | A10m, medial area 10 | DMN | 8, 58, 13 | 2.92 | 0.002 | 0.049 | 0.376 |
| | L. orbital gyrus | A14m, medial area 14 | DMN | -7, 54, -7 | 3.58 | <0.001 | 0.014 | 0.451 |
| | R. orbital gyrus | A14m, medial area 14 | DMN | 6, 47, -7 | 3.72 | <0.001 | 0.004 | 0.503 |
| | L. cingulate gyrus | A23d, dorsal area 23, posterior | DMN | -4, -39, 31 | 2.91 | 0.002 | 0.049 | 0.320 |
| | L. cingulate gyrus | A32p, pregenual area 32, anterior | DMN | -6, 34, 21 | 4.25 | <0.001 | <0.001 | 0.531 |
| | R. cingulate gyrus | A32p, pregenual area 32 | VAN | 5, 28, 27 | 2.88 | 0.002 | 0.049 | 0.392 |
| | L. cingulate gyrus | A23v, ventral area 23 | DMN | -8, -47, 10 | 3.09 | 0.002 | 0.043 | 0.396 |
| | L. cingulate gyrus | A32sg, subgenual area 32, anterior | DMN | -4, 39, -2 | 4.27 | <0.001 | <0.001 | 0.529 |
| | R. cingulate gyrus | A32sg, subgenual area 32, anterior | DMN | 5, 41, 6 | 4.03 | <0.001 | 0.003 | 0.491 |
| | L. thalamus | mPFtha | THA | -7, -12, 5 | 4.16 | <0.001 | 0.003 | 0.600 |
| R. HPC | L. sup. frontal gyrus | A9m,medial area 9 | DMN | -5, 36, 38 | 3.11 | 0.001 | 0.027 | 0.396 |
| | R. sup. frontal gyrus | A9m,medial area 9 | FPN | 6, 38, 35 | 3.04 | 0.001 | 0.029 | 0.392 |
| | L. sup. frontal gyrus | A10m, medial area 10 | DMN | -8, 56, 15 | 3.03 | 0.001 | 0.029 | 0.357 |
| | R. sup. frontal gyrus | A10m, medial area 10 | DMN | 8, 58, 13 | 2.76 | 0.003 | 0.042 | 0.347 |
| | L. orbital gyrus | A14m, medial area 14 | DMN | -7, 54, -7 | 3.24 | 0.001 | 0.027 | 0.402 |
| | R. orbital gyrus | A14m, medial area 14 | DMN | 6, 47, -7 | 3.66 | <0.001 | 0.007 | 0.498 |
| | R. cingulate gyrus | A24rv, rostroventral area 24 | FPN | 5, 22, 12 | 2.87 | 0.002 | 0.033 | 0.432 |
| | L. cingulate gyrus | A32p, pregenual area 32 | DMN | -6, 34, 21 | 4.45 | <0.001 | <0.001 | 0.540 |
| | R. cingulate gyrus | A32p, pregenual area 32 | VAN | 5, 28, 27 | 3.84 | <0.001 | 0.002 | 0.493 |
| | L. cingulate gyrus | A23v, ventral area 23 | DMN | -8, -47, 10 | 3.22 | <0.001 | 0.012 | 0.396 |
| | R. cingulate gyrus | A23v, ventral area 23 | VIS | 9, -44, 11 | 2.82 | 0.002 | 0.033 | 0.381 |
| | L. cingulate gyrus | A32sg, subgenual area 32 | DMN | -4, 39, -2 | 4.23 | <0.001 | <0.001 | 0.538 |
| | R. cingulate gyrus | A32sg, subgenual area 32 | DMN | 5, 41, 6 | 4.19 | <0.001 | <0.001 | 0.529 |
| | L. thalamus | mPFtha | THA | -7, -12, 5 | 3.85 | <0.001 | 0.002 | 0.567 |
| | R. thalamus | mPFtha | THA | 7, -11, 6 | 3.07 | <0.001 | 0.033 | 0.454 |
| | R. thalamus | rTtha, rostral temporal thalamus | THA | 3, -13, 5 | 3.27 | <0.001 | 0.019 | 0.509 |

*NMDAR: anti-N-methyl-D-aspartate receptor; HPC: hippocampus; sup: superior; mPFtha: medial prefrontal thalamus; DMN: default mode network; VAN: ventral attention network; THA: thalamus; FPN: frontoparietal network; VIS: visual network.*

**Table S2. Hippocampal BOLD pattern incongruency alterations in edges with reduced functional connectivity in NMDAR encephalitis**

| Seed region | Anatomical label | Detailed anatomical and modified cyto-architectonic description | Network label | MNI coordinates (X, Y, Z) | T | $p_{uncor}$ | $p_{FDR}$ | Effect size (Cohen's D) |
|---|---|---|---|---|---|---|---|---|
| L. HPC | L. sup. frontal gyrus | *A10m, medial area 10* | DMN | -8, 56, 15 | -2.32 | 0.012 | 0.028 | -0.357 |
| | L. orbital gyrus | *A14m, medial area 14* | DMN | -7, 54, -7 | -2.70 | 0.004 | 0.012 | -0.378 |
| | R. orbital gyrus | *A14m, medial area 14* | DMN | 6, 47, -7 | -2.98 | 0.002 | 0.012 | -0.404 |
| | L. cingulate gyrus | *A23d, dorsal area 23* | DMN | -4, -39, 31 | -2.24 | 0.014 | 0.029 | -0.331 |
| | L. cingulate gyrus | *A32p, pregenual area 32* | DMN | -6, 34, 21 | -2.07 | 0.021 | 0.033 | -0.365 |
| | L. cingulate gyrus | *A23v, ventral area 23* | DMN | -8, -47, 10 | -2.79 | 0.003 | 0.012 | -0.343 |
| | L. cingulate gyrus | *A32sg, subgenual area 32* | DMN | -4, 39, -2 | -2.09 | 0.022 | 0.033 | -0.296 |
| | L. Thalamus | *mPFtha* | THA | -7, -12, 5 | -2.74 | 0.003 | 0.012 | -0.492 |
| R. HPC | L. cingulate gyrus | *A32sg, subgenual area 32* | DMN | -4, 39, -2 | -2.53 | 0.007 | 0.037 | -0.372 |
| | L. Thalamus | *mPFtha* | THA | -7, -12, 5 | -3.13 | 0.001 | 0.011 | -0.569 |
| | R. Thalamus | *rTtha, rostral temporal thalamus* | THA | 3, -13, 5 | -3.16 | 0.001 | 0.011 | -0.522 |

*NMDAR: anti-N-methyl-D-aspartate receptor; HPC: hippocampus; sup: superior; mPFtha: medial prefrontal thalamus; DMN: default mode network; THA: thalamus.*
